# Elucidating microRNA-34a organisation within Human Argonaute-2 by DNP MAS NMR

**DOI:** 10.1101/2024.03.28.587161

**Authors:** Rubin Dasgupta, Walter Becker, Katja Petzold

**Affiliations:** Department of Medical Biochemistry and Microbiology, Uppsala University Husargatan 3, 75237 Uppsala, Sweden; Department of Medical Biochemistry and Biophysics, Karolinska Institute, 17177 Stockholm, Sweden

**Keywords:** solid-state NMR, microRNA, Dynamics Nuclear Polarisation enhancement, Human Argonaute-2, RNA dynamics

## Abstract

Understanding mRNA regulation by microRNA (miR) relies on the structural understanding of the RNA-induced silencing complex (RISC). Here, we elucidate the structural organisation of miR-34a, deregulated in various cancers, in hAgo2, effector protein in RISC, using guanosine-specific isotopic labelling and dynamic nuclear polarisation (DNP)-enhanced solid-state NMR. Homonuclear correlation experiments revealed that the non-A-form helical conformation of miR-34a increases when incorporated into hAgo2 and then bound to SIRT1 mRNA compared to the free hairpin or the free duplex formed with mRNA. Nucleotide-specific information of the of C2’- and C3’-endo sugar puckering can be obtained from the C8 – C1’ correlation with varying distributions, revealing a trapping of different confirmations upon freezing. C3’-endo puckering was predominantly observed for the seed, while C2’-endo for the central region and a mixture of both elsewhere. These observations provide insights into the molecular dynamic basis of miR-based mRNA regulation, while also providing a proof-of-concept that experiments under cryogenic conditions, e.g. at 90K, can trap and with that reveal frozen dynamic states, using methods such as (DNP-enhanced) solid-state NMR or Cryo-EM

---

MicroRNAs (miRs) are ∼22 nucleotide (nt) long RNAs that target messenger RNAs (mRNA) by binding the 3’-untranslated region (3’-UTR), and regulate these via the RNA-induced silencing complex (RISC)^[1]^. miRs regulate more than half of all mRNAs and are regularly unbalanced in cancer and other diseases^[2–5]^. The RISC core is composed of the Argonaute protein and the guide miR. Out of four isoforms, human Argonaute-2 (hAgo2, EC:3.1.26.n2, 97 kDa) displays the highest miR-based target RNA regulation activity and slicing activity^[4,6,7]^. X-ray crystal structures revealed key insights into its mechanism, however, a complete structural model of miR in hAgo2 is missing, due to the undefined electron densities of the miR central region (Figure S1)^[8–10]^ caused by the dynamic behaviour of the miR. This makes modelling this region challenging (Figure S1a – f). The crystal structures^[8–14]^ show that the seed region (nucleotide 2 – 8) of a miR is preorganised in an A-form helix with the characteristic continuous C3’-endo ribose sugar pucker (Table S1a, b Figure 1c). In absence of the mRNA target (hAgo2:miR, binary complex), the central, supplementary, and tail regions (nucleotides 9-12, 13-16 and 17-22, respectively) display non-A-form conformations characterised by the mixture of both C3’and C2’-endo ribose pucker (Table S1a). In presence of the mRNA target (ternary complex)^[11–14]^, the central region remains dynamic with undefinable electron density in the crystal structures but the supplementary region, adopts an A-form helical conformation with C3’-endo sugar pucker (Table S1b) due to base-paring with the mRNA.

**Figure 1.**
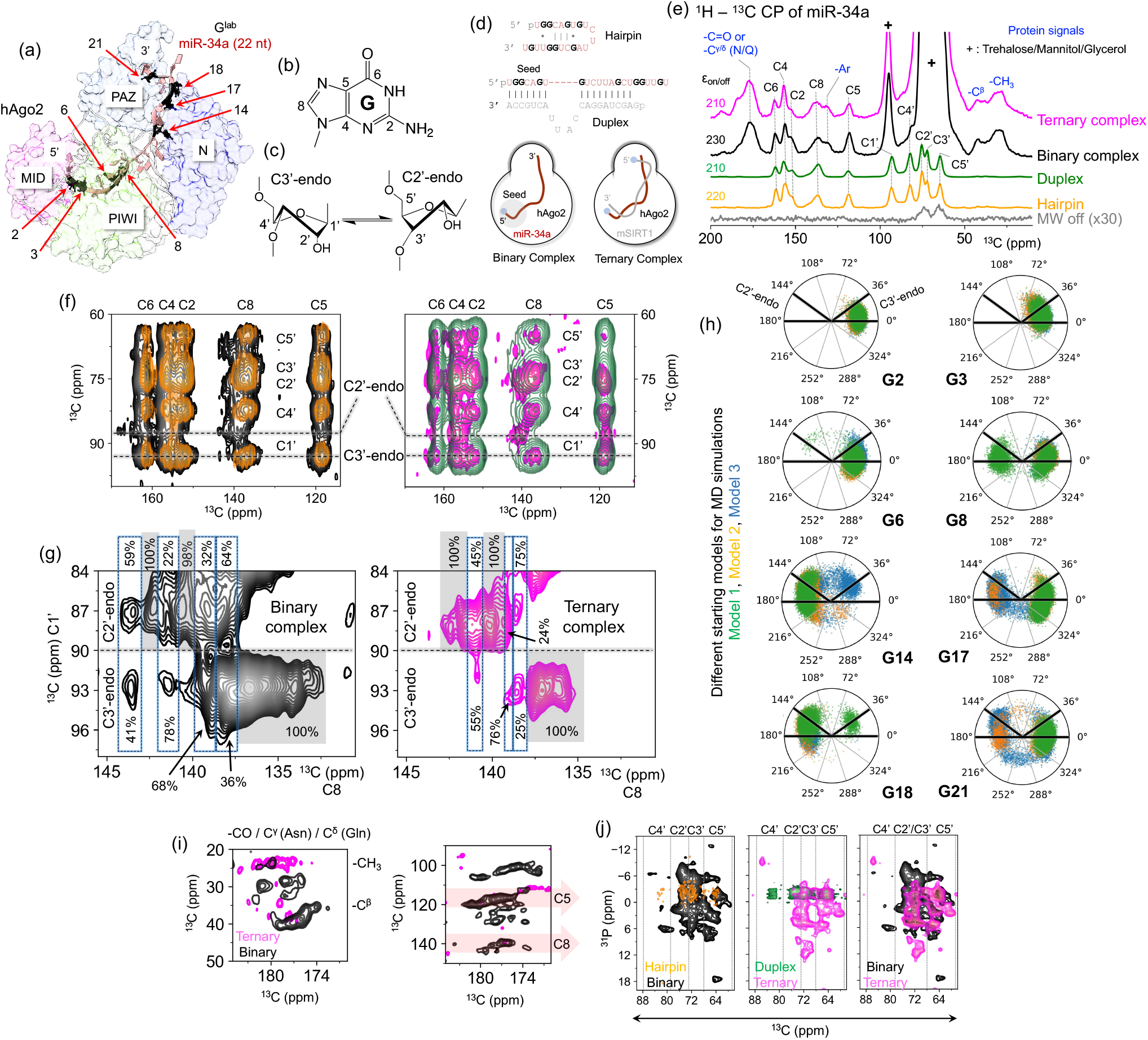
(a) Model of hAgo2:miR-34a complex based on crystal structure 4w5n^[8]^ depicting the four domains of the protein (MID pink, PIWI green, PAZ gray and N blue). Positions of ^13^C/^15^N-labeled guanosines are denoted in black and numbered. (b) Chemical structure of guanosine nucleobase with carbons numbered. (c) Depiction of equilibrium between the C3’and C2’-endo sugar puckering present in RNA. The ribose carbons are numbered in C3’-endo. (d) Overview of constructs used: top secondary structure^[1]^ of miR-34a hairpin, next miR-34a:SIRT1(mRNA) duplex, bottom schematic representation of the binary (left), and ternary (right) complexes. The seed region is shown in the duplex and binary complex. Labelled guanosines are indicated in bold. 5’-phosphate is shown as a blue circle in the binary and ternary complexes. (e) 1D ^13^C CP of G^lab^-miR-34a as a hairpin (orange), in the duplex, (green), binary (black) and ternary (magenta) complexes. Microwave (MW) off condition is shown in grey, 30X enhanced. Signal from trehalose, mannitol and glycerol in the binary and ternary complex are indicated with “+”. Signals from proteins are indicated in blue letters. Enhancement factor (ε_on/off_) relative to the glycerol signal in the MW off condition is shown for each sample. (f) 2D ^13^C-^13^C DARR displaying correlations from the nucleobase to ribose carbons overlayed with lines at 93 ppm (C3’-endo) and 87 ppm (C2’-endo) C1’ chemical shifts indicating different populations in sugar puckers accessible in all complexes and the hairpin. (g) Percentage distribution of C2’and C3’-endo conformation after deconvolution for the binary (black) and ternary (magenta) complex. The guanosines exclusively present in either of the conformations are highlighted with grey boxes, while the blue highlight shows the guanosines, with varying degree of mixed sugar puckers. (h) Ribose pucker pseudo rotation cycle indicating the distribution of each guanosine in miR-34a by MD simulation of the binary complex from three different starting models (blue, orange, and green), C2’and C3’-endo positions are denoted in G2. (i) DARR regions showing intra-hAgo2 cross-peaks (left panel) and inter hAgo2-G^lab^-miR-34a cross-peaks in binary and ternary complex (right panel), with C5 and C8 regions highlighted in red arrows. (j) Comparison of the ^13^C-^31^P TEDOR spectra from all the samples depicting the increased of spread of ^31^P resonances in binary and ternary complexes compared to hairpin and duplex. Regions of ribose carbons indicated with dashed lines.

Here, we investigate the structure and dynamics of miR-34a beyond the seed region using NMR spectroscopy. MiR-34a has been evaluated as a drug candidate against terminal liver cancer in a phase-I clinical trial with partial success^[15]^. Its structural organisation in hAgo2 however, is still unknown. miR-34a was studied (a) in its isolated hairpin form, (b) as a duplex with a shortened 21nt from 3’-UTR of the target mRNA, encoding the deacetylase sirtuin-1 (mSIRT1), (c) in a binary complex with hAgo2 (104 kDa) and (d) in a ternary complex with hAgo2 and mSIRT1 (110 kDa) (Figure 1a – d, Table S2-S4, Figure S2).

Challenges in sample purification and preparing the binary complex^[16]^, limits the available sample quantity, and conventional NMR experiments, requiring millimolar concentration, are not feasible. Therefore, Dynamic Nuclear Polarisation (DNP)-enhanced Magic Angle Spinning (MAS) solid-state NMR, where polarisation is transferred from unpaired electrons in a stable biradical to the nearby nuclei,^[17]^ is performed to provide the necessary signal enhancement. The cryo-temperatures (90 – 110 K) applied in DNP MAS NMR experiments leads to reduced resolution and increased signal overlap. Thus, ^13^C,^15^N-guanosine-labelled miR34a (G^lab^-miR-34a)^[18,19]^ representing all regions (seed nts 2, 3, 6 & 8; central nt 14; supplementary nt 17; tail nts 18 & 21)^[4,20]^, was used to mitigate resolution issues (Figure 1a – b, d, S5 – 6).

^13^C cross-polarization (CP)^[21], 13^C-^31^P Transfer Echo Double Resonance (TEDOR)^[22]^ and ^13^C-^13^C Dipolar Assisted Rotary Resonance (DARR)^[23]^ experiments (Table S5) were measured at 400.271 MHz ^1^H Larmor frequency (9.4 T), 12 kHz MAS frequency at 90 K. Samples were dissolved in DNP solvent containing 60% ^12^C,d8-glycerol, 40% H2O, 12 mM AMUPol^[24]^, 175 mM of trehalose and 329 mM of mannitol^[17,25,26]^. This led to an overall signal enhancement (εon/off, relative to the residual glycerol signal) of 220, 210, 230 and 210 for the hairpin, duplex, binary and ternary complex, respectively (Figure 1e, S3-4a, S7-8a), allowing the acquisition of homoor heteronuclear corelation spectra from 1.45 nmol G^lab^-miR-34a (60 μM in 25 μL of DNP solvent) in binary and ternary complexes respectively (Table S5).

Using the 1D ^13^C CP experiments all carbons of guanosines^[27]^ in the hairpin and duplex were detected (Figure 1e). Furthermore, for mir-34a in the binary and ternary complexes, carbonyl carbons, C^γ^ of Asn or C^δ^ of Gln (165 - 185 ppm), ring carbons of Phe, Tyr, Trp or His (120 - 130 ppm) and aliphatic -C^β/γ^ and -CH3 carbons (12 - 42 ppm)^[28]^ of the unlabelled hAgo2 were detectable (Figure 1e & i). Although, DARR spectra with 250 ms mixing time show correlations between all carbons in the hairpin and duplex, resonance assignment was hindered by extensive overlap in other regions (Figure S3c – d, S4c – d). This is likely due to signal broadening caused by freezing of multiple conformations and incomplete averaging of the chemical shift anisot_ropy_[29,30] Similar overlap is observed in the binary and ternary complexes. Interestingly, the C1’ resonances are split into two regions (Figure 1e-g, S7c-e and S8c-e) and signals between 90.4 – 96.3 ppm can be attributed to C3’-endo sugar pucker with A-form helical conformation, while resonances between 86.6 – 90.0 ppm are associated to the C2’-endo sugar pucker with non A-form geometry^[28,31–35]^. The positive projection of these regions from each of the systems studied shows that there is a 14.0 ± 0.2% and 11.3 ± 0.3% population of C2’-endo sugar pucker in the hairpin and duplex, respectively, based on Gaussian fits to the line shape. This increases to 34.4 ± 1% and 43.8 ± 1% for the binary and ternary complex, respectively, (Figure S9, Table S6-S9) and indicates that the μs timescale dynamics of the sugar puckering^[34]^ can be frozen into their respective ensembles and trapped for detection under DNP conditions. For the hairpin and duplex, the distribution of the C2’and C3’-endo pucker in guanosines is consistent with the population distribution reported earlier for helical nucleotides at room temperature (14%:88% for C2’:C3’-endo)^[1,34]^.

Evidently, the protein induced changes the population distributions (Figure S9) of the sugar puckers. Crystal structures of binary complexes (Table 5a) suggest that nucleotides 2, 3, 6, 8 and 21 are in the C3’-endo conformation while nucleotides 14, 17 and 18 are in the C2’-endo conformation leading to a theoretical distribution of 62.5% (C3’-endo):37.5% (C2’-endo). This agrees with the fit of the regions in the DARR spectrum (Figure S9c) that yields a distribution of 65.6 ± 1% (C3’-endo):34.4 ± 1% (C2’-endo). In the ternary complex, the proportion of the C2’-endo increases. All the reported crystal structures (Table 5b) of the ternary complex shows that the nucleotides 2 – 8 of the guide RNA remain in the C3’-endo conformation, thereby ruling out the possibility that these nucleotides are responsible for the increased proportion of C2’-endo conformation. Crystal structures 6N4O, 6NIT, 6MFR and 6MDZ^[12,13]^ show that the central and the supplementary region display C3’-endo conformation in presence of the mRNA target (Table 5b). This contrasts the observed fit of the DARR spectrum (Figure S9) from the ternary complex and is likely due to the RNA duplex release^[12,13,36]^ from hAgo2. Little to no interaction of the RNA duplex with the PAZ domain of the protein is observed in the crystal structures 6MFR, 6MDZ and 6NIT^[12]^ (Figure S9b). Optimisation of the experimental conditions in the present study mitigates duplex release from the protein when using a shortened 21nt target mRNA as reported previously^[13,36,37]^. This is evident from the C3’ and C2’-endo conformation distribution in the ternary complex (Figure 1g), which is different from the duplex sample.

In addition to this overall puckering distribution, the high-resolution in the C8 – C1’ correlation region allowed nucleotide-specific analysis of the puckering from the binary and ternary complexes. Gaussian peak fitting revealed 12 and 11 peaks for the whole C8 region of the 8 guanosines for the binary and ternary complexes, respectively. These additional resonances can originate from the changes around the glycosidic bond (χ-angle) to which the C8 chemical shift is sensitive.

It is observed that the guanosine specific dynamic sampling of C2’and C3’endo sugar puckering is frozen. In the binary complex, 7 resonances are exclusively in either conformation (4 C3’and 3 C2’-endo), while 4 resonances showed distribution of 64:36%, 32:68%, 22:78% and 59:41% (error of < 1%) for C2’:C3’endo sugar puckering (Figure 1g and S10, Table S10). The ternary complex is less conformationally diverse with 8 resonances remaining in a single conformation (4 C3’- and 4 C2’-endo) and while 3 resonances have two conformations of 75:25%, 24:76% and 45:55% (error of < 1%), respectively (Figure 1g and S10, Table S11). The varied distributions indicate that each nucleotide can have different tendency to populate C3’-(A-form) or C2’-endo (non-A-form) conformation, implying that cooling the sample does not promote the selection of a single conformation or a coalescing to the lowest energy conformation. Additionally, the fact that the ternary complex spectrum does not collapse into the duplex spectrum (Figure 1f, S9) shows that the duplex release is suppressed. To identify individual guanosines responsible for the different puckering signals, miR-34a was modelled into hAgo2 using crystal structure 4W5N^[8]^ for the binary complex and simulated from three different starting structures for 400 ns each amounting to a total of 1.2 μs. The puckering distribution of the guanosines G2, G3 and G6 indicated an exclusive in C3’-endo, while G14 and G18 remained in C2’-endo, and G8, G17, and G21 experience mixed conformations (Figure 1h). Mapping this distribution onto the DARR region of C8 – C1’ correlation from the binary complex it can be hypothesized that resonances between 132.99 and 136.83 ppm with C3’-endo conformation belongs to G2/3/6. This is consistent with reported crystal structures (Table 5a), where the seed region is shown to be in C3’-endo conformation. Signals between 139.78 and 140.63 ppm would then belong to G14/18, while the signals with a mixture of the C2’- and C3’-endo conformation could be G8/17/21. A splitting of resonances due to trapped *anti* / *syn* conformations is likely to be found for G8, G14, G18 and G21 (Figure S11).

Interestingly, cross-peaks between the unlabelled hAgo2 (180 – 174 ppm) and G^lab^-miR-34a (145 – 112 ppm) as well as intra-protein cross peaks are readily observed in the binary complex (Figure 1i); enabled by a DARR mixing time of 250 ms that can yield correlations between carbons of up to 6 Å in distance^[38]^. The MD simulation trajectories were further analysed, by (a) selecting amino-acids withing a 6 Å zone around the guanosines of miR-34a and (b) comparing their carbon chemical shift with the Biological Magnetic Resonance Data Bank (BMRB). This allowed the identification of DARR correlations to cross peaks likely arising from Cys66 CO – G14C8, Asn551 C^γ^ – G2C5/C8, and Thr559 CO – G2C5/C8 (Figure 1i and S12). These cross-peaks, however, were not observed in the hAgo2:G^lab^-miR-34a:SIRT1 ternary complex (Figure 1i). Since there is no duplex release (Figure 1j middle, details above), this observation can be due to structural rearrangement imposed by the presence of the mRNA target where the nucleobase of miR-34a gets oriented to form base pair supported by the chemical shift perturbations in the C8 to C5/6/4/2 correlations (Figure S13). This leads to an increased distance between the nucleobase carbons and the respective amino acids hence decreasing the cross-peak intensity beyond detection.

Additional information on the structural organisation of miR34a in hAgo2 can be obtained from ^31^P chemical shifts spanning the backbone of the RNA. The 2D ^31^P-^13^C TEDOR spectra reveals that the ^31^P resonances in the binary complex have a large chemical shift dispersion of ∼16 ppm compared to ∼6 ppm in the hairpin (Figure 1j, S7g-S8f). The BMRB shows a spread of ∼ 8 ppm (-6 to 2 ppm from 74 entries as of February 2024 where two entries represent protein-RNA complex) in the ^31^P chemical shift for guanosines in RNA, which do not explain the current observation. Many factors can affect the ^31^P chemical shift variation^[39–44]^. A-form RNA changes do alone not explain the increased chemical shift spread, however, variations in backbone torsion angles or O-P-O bond angles might contribute to the increased chemical shift spread. Additionally, the observed chemical shift dispersion can be a combination electrostatic potential and hydrophobicity in the binding pocket of protein (Figure S14). Furthermore, the similarity of the ^31^P chemical shift dispersion between the binary and ternary complexes (Figure 1j) suggests that the RNA duplex is stable within hAgo2 and the duplex release^[13,45–47]^, even partially, is supressed, consistent with the above observations.

In conclusion, using DNP MAS NMR, with a signal enhancement of ∼ 200 enabled the structural characterisation of the 104 kDa binary and 110 kDa ternary complex. Due to the high signal enhancement, correlations between G^lab^-miR-34a and unlabelled hAgo2 were detected and yielded detailed information about the conformational properties of miR-34a in the RISC. ^31^P chemical shift perturbations and ribose pucker distribution indicate that the duplex release phenomenon is supressed in the ternary complex. We were able to detect conformational ensembles being trapped in the frozen states with varying distribution of the sugar pucker – C2’-vs C3’-endo, indicating that DNP-enhanced solid-state NMR can monitor room temperature conformational heterogeneity under frozen conditions. Similar observations of frozen multiple conformations were made in DNP studies of proteins both *in vitro* and in cell^[48]^, and complementary to that, in cryo-EM studies^[49,50]^ For large nucleic acids protein complexes the dynamic information so far has been extrapolated from the local disorder in crystal structure (B-factor) or the lack of defined electron density therein^[8,9,13]^. Recent cryo-EM structure of plant Argonaute has shown two different conformations of the RNA duplex in the ternary complex, though, due to low local resolution detailed puckering information could not be obtained^[51]^. The present study provides the first observation of the different puckering preference of individual nucleotides in various regions of miR-34a including the dynamic central region, which has not been observed in the crystal structures. It also highlights the capability of DNP MAS NMR to trap room temperature dynamic states and thereby gain structural information on low-populated states. Moreover, the presence of a 21nt SIRT1 target mRNA changes the conformational preferences and decreases the non-helicity of miR-34a by ∼10 %. This is expected to differ for other targets of miR-34a, hence opening the possibilities to further study its conformation within hAgo2 in the presence of multiple targets, ultimately leading to a comprehensive picture of miR-based target discrimination and regulation.

## Acknowledgements

The Protein Science Facility (PSF) at Karolinska Institute is acknowledged to produce T7 polymerase, RNase H, and TEV protease. The Protein Expression and Characterization platform, SciLifeLab, Solna is thanked for providing the facility to produce hAgo2 in SF9 Insect cells. SwedNMR national NMR facility at Gothenburg University is acknowledged for providing the DNP MAS NMR instrument. R.D. acknowledges funding from European Union’s Horizon 2020 research and innovation programme under the Marie Skłodowska-Curie grant agreement No. 101067627, project: ECONOMICS. K.P. acknowledges funding from Wallenberg Academy Fellow (KAW 2019, 0227), project grant from the Knut och Alice Wallenberg foundation (KAW 2016.0087), Cancerfonden (CAN 2018/715 & 21 1770 Pj-BF 1), KI consolidator grant (2-2111/2019) and Karolinska Institute for the help with the purchase of our 600 MHz NMR. The computations were enabled by resources provided by the National Academic Infrastructure for Supercomputing in Sweden (NAISS), partially funded by the Swedish Research Council through grant agreement no. 2022-06725. We thank the Petzold lab for discussion.

