## Supporting Information for "Elucidating microRNA-34a organisation within Human Argonaute-2 by DNP MAS NMR"

### Supporting Materials and methods

*RNA production and purification*

$G^{lab}$-miR-34*a* and 21nt-SIRT1 mRNA were produced using *in vitro* transcription (IVT)[1]. The DNA template and the associated T7 promoter for SIRT1 were purchased from Integrated DNA Technologies (IDT). 1 ml of 20 μM solution of the DNA template and T7 promoter were annealed by heating it at 95°C for 5 min and then incubated at room temperature for an hour. This was used for a 10 ml IVT reaction (100 mM Tris-glutamate pH 8.0, 15 mM $Mg(OAc)_2$, 10 mM DTT, 10% DMSO, 25 mM Spermidine, 5 mM GMP, 3 mM of each NTP, 0.4 μM DNA and 0.22 mg/ml S43Y T7 polymerase[2]). The mixture was incubated at 37°C overnight. The reaction was quenched with 500 mM EDTA pH 8.0 to a final concentration of 20 mM. The solution was buffer-exchanged to water and concentrated to 2ml. Equal volume of High Dye pH 8.0 (0.25 mg/ml Bromophenol blue, 1X TBE pH 8.0, 50 mM EDTA dissolved in formamide) was added and stored at -20°C for gel purification. SIRT1 was purified using 4 15% denaturing PAGE casted on a 16.5 (w) cm X 38.7 (h) cm plate with a 1.5 mm spacer and comb (CBS scientific). The gels were run at 80W for 3 h 30 min and the gel was imaged using UV shadowing. The most intense band was excised and incubated with 3 ml TEN buffer (10 mM Tris pH 8.0, 1 mM EDTA and 300 mM NaCl) at 37°C for 2 to 3 hrs. The supernatant was collected by filtering the gel pieces through a 0.2 μM cellulose acetate syringe filter. The supernatant (warm extract) was stored at -20°C for further processing. The gel pieces were incubated again with 3ml of TEN buffer at 4°C overnight. The supernatant (cold extract) was collected and stored at -20°C. The warm and cold extract from all 4 gels were pooled into their respective fractions and the purified RNA was ethanol precipitated. The two extracts were processed separately. For ethanol precipitation, 3x volume of chilled 100% ethanol was added to the extracts and incubated on dry ice for an hour. This solution was centrifuged at 4°C with 13000 rpm for 1 hour. The supernatant was discarded carefully without disturbing the RNA pellet. 300 μL of 70% chilled ethanol was added to the pellet slowly via the side of the tube to not disturb the pellet. This was centrifuged at 4°C with 13000 rpm for 30 min. The supernatant was carefully discarded. The pellet was dried under vacuum for 30 min and then dissolved in 1ml of RNase free water. The solution from both the extracts were pooled together. The purity was checked using 20% denaturing PAGE. The total yield of SIRT1 was 96 nmoles.

$G^{lab}$-miR-34a was prepared by IVT of a linearized DNA template with 26 tandem repeats followed by RNaseH cleavage of the transcript[3]. The pUC19 plasmid containing the tandem repeats of miR-34a was propagated in chemical competent DH5α *E.coli* and purified using a miniprep kit from Thermofischer scientific. 20 µg/ml of plasmid was linearized using 150 units of BamH1-HF (New England BioLabs Inc.) for an hour at 37°C. The reaction was stopped by heating the solution at 80°C for 10 min and then stored at -20°C. 10 mL IVT reaction (100 mM Tris-glutamate pH 8.0, 25 mM $Mg(OAc)_2$, 10 mM DTT, 25 mM Spermidine, 5 mM GMP, 3mM of each NTP, 20 µg of plasmid, 20 mM chimera, 25 µg/mL RNaseH and 0.22 mg/ml S43Y T7 polymerase[2]) with $^{13}C/^{15}N$ labeled guanosine triphosphate (Sigma-Aldrich 645680-25MG) was incubated at 37°C overnight. The reaction was processed and purified like the SIRT1 with a total yield of 162 nmol. The purity was checked using 20% denaturing PAGE.

*Bacmid and baculovirus production*[4]

The gene construct was created by cloning the full-length human argonaute-2 protein (hAGO2) into pFastBac dual expression vector (ThermoFisher scientific, catalog number: 10712024) between the BamHI and EcoRI restriction sites using the standard double digestion and ligation protocol as given by NEB. The resulting hAgo2-pFastBac construct contained a 6xHis-tag-TEV cleavage site at the N-terminus of the cloned gene. The gene is controlled by the polyhedrin promoter for expression. Recombinant Bacmid was produced by transforming 100 ng of hAGO2-pFastBac construct into electrocompetent DH10EmBacY cells (Geneva Biotech). Multiple rounds of phenotype verification were performed using blue/white screening, bacmid isolation, and PCR analysis according to the "Bac-to-Bac TOPO Expression System" user manual (Thermofisher, Version A,15 December 2008 A10606) to confirm hAGO2 containing recombinant Bacmid. Initial transfection into SF9 insect cells (P0-hAgo2) was done by packing the recombinant Bacmid in cationic lipids (Cellfectin II reagent, Thermofisher) and exposing to fresh 8 x $10^5$ Sf9 cells/well (1.5 x $10^6$ cells/ml) for 4 hours at 27°C. Two consecutive rounds of baculovirus stock amplification were performed, starting from P0-hAgo2 and increasing the stock volumes from 25 ml (P1-hAgo2) to 300 ml (P2-hAgo2). The final 300 ml P2-hAgo2 baculovirus stock was filtered and supplemented with FBS (2% final conc.) for long-term storage. 1.5 ml aliquot of P2-hAgo2 was used for western blot analysis using Anti-AGO2 mouse monoclonal antibody (clone: 2E12-1C9, VWR art. number: USBI126228) and Anti-Argonaute-2 monoclonal antibody (EPR10411, abcam identifier: ab186733) binding to either N-domain (N-terminal, Linker-1, and PAZ) or C-domain (Linker-2, MID and PIWI) of hAgo2 respectively to confirm its expression after 72 hrs. The P2-hAGO2 was used for large-scale production.

*Insect cell protein expression*

The protein expression was initiated by adding 10 mL of P2-hAGO2 into 1 L of SF-900™ II SFM (ThermoFischer scientific, catalog number: 10902096) containing Sf9 cells at a confluency of 1.75 x $10^6$ cells/mL (passage 14). A 3 L Erlenmeyer flask (Corning®, Fernbach design, product number: 431252) per liter of the cells was used for the production. The cells were grown in an orbital shaker at 120 rpm, 27°C for 48 h without $CO_2$. Throughout the overexpression period, live cell numbers and protein expression levels were monitored every 24 hours using trypan blue staining (ThermoFischer scientific, catalog number: 15250061). The expression was also monitored by the Green Fluorescent Protein (GFP) levels. The cells were observed using CelenaS, Logos Biosystems cell imager. After 48 h of overexpression, 6 L cell culture was centrifuged at 900 RCF, washed with phosphate buffer saline solution at 1300 RCF and the cell pellet was collected. The total wet cell mass obtained was 45 g. This was stored at -80°C for future protein purification.

*Protein purification*

22 g of cell pellet was resolved in 25 mL of pre-cooled $Ni^{2+}$-affinity equilibration buffer (50 mM Tris-HCl pH 8.0, 300 mM NaCl, and 1 mM TCEP). This was supplemented with 1 mL of 25x EDTA-free protease inhibitor cocktail (Merck, product number: 11873580001). The cells were lysed by multiple freeze thaw cycles using dry ice and water bath at room temperature. The final cell lysis was done using sonication for 4 min at 80% amplitude with 5 sec on and 5 sec off cycles. This lysate was centrifuged for 2 hrs at 4°C and 30,000 RCF. The supernatant was collected and filtered through a 0.22 µm PES membrane (Millipore Express® PLUS) using a sterile vacuum filtration system. The filtered supernatant was loaded onto a pre-equilibrated 5 mL HisTrap-$Ni^{2+}$ column (Cytiva, HisTrap HP) and washed with 3 column volumes (CV) of equilibration buffer. Elution was done by a linear gradient of 20 CV for 100 min using the elution buffer (equilibration buffer + 300 mM Imidazole). Fractions were run on a SDS PAGE gel and the fraction with protein was pooled together which resulted in a total volume of 15 mL. This was concentrated to 5 mL using a 30 kDa cut-off Amicon filter (Merck). The 5 mL solution was supplemented with 600 µL of crude $G^{lab}$-miR-34a obtained after concentrating 3 mL overnight IVT reaction via a 3 kDa cut-off Amicon filter (Merck). This mixture was

incubated for 1.5 h at 37°C water bath to load the hAgo2 with $G^{lab}$-miR-34a. After incubation, the solution was supplemented with 100 µL of TEV protease (produced by Protein Science Facility, Karolinska Institute, Solna, Sweden). This solution was dialyzed using 3 kDa cut-off dialysis bag (Spectrum™) against 2 L of $Ni^{2+}$-affinity equilibration buffer overnight at 4°C. The precipitate was removed by centrifuging at 4°C, 4000 RCF for 30 min. The supernatant was concentrated to 1.5 mL using a 30 kDa cut-off Amicon filter (Merck). This concentrated solution was loaded onto a pre-equilibrated (20 mM HEPES pH 7.5, 100 mM KCl, 1 mM TCEP) size exclusion column (Cytiva, Superdex 200 increase 10/300 GL). The fractions with the protein were identified on a SDS PAGE gel, pooled together, and concentrated to the final volume of 1 mL using a 30 kDa cut-off Amicon filter (Merck).

*Protein quantity and quality estimation*

The protein concentration was estimated by Bradford-based assay (ThermoFischer scientific, Pierce™ Detergent Compatible Bradford Assay Kit) in a clear 96 well plate (F-bottom, Corning). The analysis was performed as described in the user manual for microplate procedure from ThermoFischer scientific. The absorption was measured at 595 nm on a Varioskan LUX multimode microplate reader (Thermofisher). The $G^{lab}$-miR-34a loaded onto hAgo2 was quantified by Northern blot analysis. In brief, 3 µL of hAgo2:miR-34a from the 1 mL stock and different amounts of pure miR-34a (0.5, 1, 3, 5, 8, 10 and 22 pmol) was mixed with the loading dye (per 50 mL: 8 mg of bromophenol blue, 931 mg $Na_2EDTA.2H_2O$, 50 mL formamide) in a 1:5 ratio. These were completely loaded on a 20% denaturing Urea-PAGE gel and ran for 1 h at 350 V. This gel was blotted onto a nylon membrane (Immobilon-Ny+ Membrane, 0.45 µm, Millipore) for 2.5 h using a Mini Trans-Blot cell (BioRad). The miR-34a was crosslinked on the membrane with 12 mL EDC (1-Ethyl-3-[3-dimethylaminopropyl]carbodiimide, Sigma) reaction mixture for (0.377 g EDC in aqueous 130 mM methyl imidazole solution, pH 8) 1 h at 60°C in a wet environment. The cross-linked membrane was incubated with 1 nmol of complementary Cy3-labeled miR-34a probe in miliQ water overnight at 37°C in an orbital shaker with 120 rpm. The miR-34a was quantified by monitoring the Cy3 signal and intensity analysis using Image Lab software (BioRad). Splicing assay was performed according to previously reported protocols[5–7]. The equations to estimate the amount of total protein (equation 1) and miR-34a loaded in hAgo2 (equation 2) from Bradford assay and northern blot respectively are as follows:

$$Protein\ (\mu g/mL) = 1000 * \frac{Absorbance\ 595\ nm\ (sample)\ - 0.63}{0.6} \quad (1)$$

$$RNA\ (pmol) = \frac{Intensity\ (sample)\ - 1e^{6}}{7.34e^{5}} \quad (2)$$

*Sample preparation for NMR measurements*

The hairpin $G^{lab}$-miR was prepared by folding the sample in the NMR buffer (15 mM NaPi pH 6.5, 25 mM NaCl and 0.1 mM EDTA) by heating at 95°C for 5 mins followed by snap cooling on ice for 20 mins. $G^{lab}$-miR-34a:SIRT1 duplex was prepared by mixing $G^{lab}$-miR-34a and SIRT1 in 1:1 ratio (80 nmol each) in NMR buffer and heating the solution for 5 mins at 95°C followed by slow cooling to 25°C for a total time of 3 h. For solution NMR experiments, 80 nmol of the hairpin were dissolved in NMR buffer containing 6% $D_2O$. These samples were lyophilized and re-measured under the same condition to control for effects of lyophilization. For DNP experiments these samples were recovered, buffer-exchanged to nuclease-free water using 3 kDa cut-off Amicon filter and supplemented with 9 µL of NMR buffer. This mixture was lyophilized and stored at -20°C.

750 µL of hAgo2:$G^{lab}$-miR-34a (binary) complex was buffer-exchanged to nuclease-free water using 50 kDa cut-off Amicon filter and supplemented with 9 µL of NMR buffer, and 175 mM of trehalose (Fischer Scientific 15477189) and 329 mM of mannitol (Fischer Scientific 10725531) as a cryoprotectant. This mixture was flash-frozen in liquid nitrogen, lyophilized overnight and stored at -20°C. The hAgo2:$G^{lab}$-miR-34a:SIRT1 (ternary) complex was formed by first mixing 750 µL of the binary complex (buffer exchanged as mentioned above) and 1 µL RNase Inhibitor (Merck, product number: 3335399001). The solution was incubated at 25°C for 15 min with shaking at 300 rpm. 1.5 nmol of SIRT1 target RNA was added to this solution and incubated for 1.5 h. at 37°C with shaking at 300 rpm. This solution was supplemented with 175 mM of trehalose and 329 mM of mannitol flash frozen with liquid $N_2$, lyophilized, and stored at -20°C.

For dynamic nuclear polarization (DNP) magic angle spinning (MAS) NMR, the lyophilized powder was resuspended in 9 µL nuclease-free water containing 2 mM $MgCl_2$. This 9 µL solution was added to 15 µL of $^{12}C$-$d_8$-glycerol (Cambridge Isotope Laboratory CDLM-8660-1) containing 12 mM AMUPol (1 µL from a stock of 300 mM solution prepared in nuclease-free water). The total volume of 25 µL was transferred into a 3.2 mm sapphire rotor (Cortec Net, product number: B6939). The sample within the rotor was sealed with a soft silicone plug (Cortec Net, product number: H176385) before sealing with $ZrO_2$ drive cap.

*NMR measurements and data analysis*

Solution state NMR was performed in a 600.16 MHz (14.1 T) magnet equipped with QCI-P H/C/N/P cryoprobe. The 1D $^{1}H$ spectra were acquired using a 90° pulse followed by excitation-sculpting water suppression with $^{13}C$ and $^{15}N$ decoupling using grap4[8] during acquisition. 32 scans were collected, and spectra were processed in Topspin 3.6.3.

DNP MAS NMR measurements were carried out at the Swedish NMR center hosted at Göteborg University, Sweden. Measurements were performed at the 400.271 MHz (9.4 T) magnet equipped with a 3.2 mm wide bore MAS DNP probe with X/Y/H channels. The X channel was tuned to $^{13}C$ and Y to $^{31}P$. The microwave (MW) source was a 275 GHz gyrotron. All the experiments were done at 90 K and MAS frequency of 12 kHz. $^{1}H$ – $^{13}C$ CP MAS experiment was done with MW on/off condition to observe the enhancement factor $\varepsilon_{on/off}$. The total intensity of the region between 200 and 10 ppm region was used to estimate the enhancement factor. The duration of the 90° $^{1}H$ excitation pulse was 2.5 μs at 100 kHz. CP duration was 2.25 ms with a rectangular pulse on $^{13}C$ and a Ramp 90-100 pulse on $^{1}H$ channel respectively. 16 scans were acquired with the acquisition time and recovery delay set to 17 ms, and 5 s respectively. The $^{13}C$ and $^{1}H$ carrier frequencies were -50.662 ppm and -129.316 ppm respectively. $^{1}H$ decoupling during acquisition was done using 52 kHz spinal64. The $^{13}C$ – $^{13}C$ dipolar assisted rotational resonance (DARR)[9] spectra were collected with the same CP parameters with DARR mixing time of 250 ms with $^{1}H$ B1 field set to the MAS frequency (12 kHz). The 90° $^{13}C$ pulse length was 3.4 μs at 73.5 kHz. $^{13}C$ - $^{31}P$ transfer echo double resonance (TEDOR)[10] experiments were performed with CP transfer from $^{1}H$ to $^{13}C$ for detection and included rotor synchronized 180° pulses on the $^{31}P$ channel during the mixing period. The 180° $^{31}P$ pulse length was 6.6 μs at 75.8 kHz with the carrier frequency set to -136.702 ppm. A 50 kHz $^{1}H$ decoupling was included during the mixing and acquisition period. 1D $^{13}C$ - $^{31}P$ TEDOR experiments were performed with 32 scans with 50 kHz spinal64 decoupling during acquisition. In the 2D experiment the t1 increment delay was rotor synchronized such that the spectral window of the indirect dimension ($^{31}P$) was half of the MAS frequency. Other important experimental parameters for DARR and TEDOR are tabulated in Table S3 for different samples. The $^{13}C$ and $^{31}P$ spectra were referenced to the room temperature resonance of glycerol and phosphate buffer pH 6.5 respectively. The data were processed using Topspin 3.6.3. The data was plotted and the gaussian fits were performed using an in-house Python 3.10 script.

*Molecular dynamics (MD) simulation*

The G$^{lab}$-miR-34a:hAgo2 complex was modeled using PDB 4W5N[6]. The missing segment in the protein was modeled using the SWISS-MODEL webpage (https://swissmodel.expasy.org/)[11]. The missing segments in the RNA were modeled manually by constraining the already present nucleotide using UCSF Chimera. The sequence of guide-RNA was mutated to miR-34a using the “swapna” command in UCSF Chimera. Amber14sb-OL15 force field (https://fch.upol.cz/ff_ol/gromacs.php) was used for the simulation. The parameters for the 5’ terminal phosphate was set to values as implemented in the CHARMM36 force field. GROMACS 2023 suite was used for the all-atom MD simulation using TIP3p water model in a cubic box with dimension 1 x 1 x 1 nm. The NaCl concentration was set to 125 mM and the charge was neutralized using excess chloride ion. Minimization was performed via the steepest descent gradient method for 50.000 steps to a minimum force constant < 1000 kJ mol$^{-1}$ nm$^{-1}$. The system was equilibrated for a total of 200 ps at 300 K using velocity rescale scheme and 1 bar pressure using Parrinello-Rahman barostat. Verlet nonbonded cut-off scheme with grid neighbor search and 1.0 nm cut-off for van der Waals interaction with energy and pressure dispersion correction was used. Particle Mesh Ewald with fourth order cubic interpolation was used for Coulomb interactions. All bonds were constrained using the LINCS algorithm. During MD run 2 fs integration step was used and trajectory was saved every 10 ps. Three production runs with different starting structures were performed for 400 ns each amounting to a total time of 1.2 µs. The trajectories were analyzed using UCSF Chimera and in-house Python 3.08 script using MDtraj[12] and Barnaba[13] plugins for sugar puckering. The coordinates of the system, are provided in the GitHub link below:
https://github.com/PetzoldLab/DNP_MAS_NMR_Glab-miR-34a

The three starting structures were simulated to have a representative conformational sampling of different sugar puckering and pseudorotation angles of the guanosines in the binary complex. The correlation of the pseudo-rotation angles and glycosidic dihedral angles (Figure S11) shows that the seed region (G2/3/6) has a stable conformational sampling, while the regions beyond the seed are sampled in many different combinations of sugar puckering and glycosidic dihedral angle, which are possibly not fully explored.

**Table S1a**: Ribose puckering of each nucleotide of miR in hAgo2 in the binary complex from the reported crystal structures. The overall resolution of the crystal structure is given in parenthesis. The definition of the glycosidic dihedral angle follows reference [14]

| Position | Nucleotide parameters | | | | | | | | |
|---|---|---|---|---|---|---|---|---|---|
| | 4W5N (2.90 Å) [6] | | | 4OLA (2.30 Å) [5] | | | 4F3T (2.25 Å) [15] | | |
| | Sugar pucker | Glycosidic bond | χ (°) | Sugar pucker | Glycosidic bond | χ (°) | Sugar pucker | Glycosidic bond | χ (°) |
| 1 | C2' | high-*anti* | -120.35 | C2' | high-*anti* | -102.21 | C2' | -122.057 | high-*anti* |
| 2 | C3' | *anti* | -156.85 | C3' | *anti* | -157.45 | C3' | -158.179 | *anti* |
| 3 | C3' | *anti* | -156.84 | C3' | *anti* | -151.30 | C3' | -146.97 | *anti* |
| 4 | C3' | *anti* | -145.54 | C3' | *anti* | -140.40 | C3' | -135.102 | high-*anti* |
| 5 | C3' | *anti* | -156.43 | C3' | *anti* | -142.39 | C3' | -132.498 | high-*anti* |
| 6 | C3' | *anti* | -162.42 | C3' | *anti* | -143.90 | C3' | -126.068 | high-*anti* |
| 7 | C3' | *anti* | -156.39 | C3' | *anti* | -158.97 | C3' | -165.925 | *anti* |
| 8 | | | | | | | C3' | -168.797 | *anti* |
| 9 | | | | | | | C3' | -143.783 | *anti* |
| 10 | | | | | | | C3' | -168.912 | *anti* |
| 11 | | | | | | | | | |
| 12 | C3' | *syn* | 54.106 | | | | | | |
| 13 | C2' | high-*anti* | -102.75 | | | | | | |
| 14 | C2' | high-*anti* | -83.016 | | | | | | |
| 15 | C2' | high-*anti* | -102.75 | | | | | | |
| 16 | C2' | *syn* | 87.19 | | | | | | |
| 17 | C2' | high-*anti* | -108.20 | | | | C3' | -162.32 | *anti* |
| 18 | C2' | high-*anti* | -118.80 | | | | C2' | 20.59 | *syn* |
| 19 | C2' | *anti* | -145.85 | | | | C3' | 75.854 | *syn* |
| 20 | C2' | high-*anti* | -107.90 | | | | C3' | -131.798 | high-*anti* |

| | | | | | | |
|---|---|---|---|---|---|---|
| 21 | C3' | *anti* | -145.97 | C3' | *anti* | -153.07 |

**Table 1b:** Ribose puckering of each nucleotide of miR in hAgo2 in the ternary complex from the reported crystal structures. The overall resolution of the crystal structure is given in parenthesis. Positions denoted as bold and underlined indicate nucleotides that are interacting with the target mRNA. The definition of the glycosidic dihedral angle follows reference [14]

| Position | Nucleotide parameters for guide miR | | | | | | | | |
|---|---|---|---|---|---|---|---|---|---|
| | 4W5O (1.80 Å) [6] | | | 4W5Q (3.10 Å) [6] | | | 4Z4C (2.30 Å) [16] | | |
| | Sugar pucker | Glycosidic bond | χ (°) | Sugar pucker | Glycosidic bond | χ (°) | Sugar pucker | Glycosidic bond | χ (°) |
| 1 | C2' | high-*anti* | -124.52 | C1'-exo | high-*anti* | -126.62 | C2' | -126.12 | high-*anti* |
| 2 | **C3'** | ***anti*** | **-164.73** | **C3'** | ***anti*** | **-166.57** | **C3'** | **-169.31** | ***anti*** |
| 3 | **C3'** | ***anti*** | **-158.03** | **C3'** | ***anti*** | **-143.67** | **C3'** | **-159.47** | ***anti*** |
| 4 | **C3'** | ***anti*** | **-154.73** | **C3'** | ***anti*** | **-154.47** | **C3'** | **-157.38** | ***anti*** |
| 5 | **C3'** | ***anti*** | **-154.34** | **C3'** | ***anti*** | **-156.89** | **C3'** | **-161.68** | ***anti*** |
| 6 | **C3'** | ***anti*** | **-158.54** | **C3'** | ***anti*** | **-172.23** | **C3'** | **-155.59** | ***anti*** |
| 7 | **C3'** | ***anti*** | **-159.86** | **C3'** | ***anti*** | **-156.34** | **C3'** | **-164.26** | ***anti*** |
| 8 | **C3'** | ***anti*** | **-154.68** | **C2'-exo** | ***anti*** | **-128.21** | **C3'** | **-157.09** | ***anti*** |
| 9 | **C3'** | ***anti*** | **-150.25** | C3' | *anti* | -147.85 | **C3'** | **-154.08** | ***anti*** |
| 10 | C2' | high-*anti* | -102.06 | C1'-exo | high-*anti* | -112.28 | C2' | -102.75 | high-*anti* |
| 11 | C3' | *anti* | -165.19 | C3' | *anti* | 175.42 | C3' | -166.06 | *anti* |
| 12 | C3' | *anti* | -164.23 | C3' | *anti* | -160.18 | C3' | -162.88 | *anti* |
| 13 | C3' | *anti* | -164.29 | C3' | *anti* | -166.33 | C3' | -165.87 | *anti* |
| 14 | C3' | *anti* | -162.43 | C3' | *anti* | -157.33 | C3' | -160.09 | *anti* |
| 15 | C3' | *anti* | -152.01 | C3' | *anti* | -150.37 | C3' | -154.97 | *anti* |
| 16 | C2' | high-*anti* | -102.91 | C2' | high-*anti* | -102.36 | C2' | -79.80 | high-*anti* |
| 17 | C3' | *anti* | -160.00 | C3' | *anti* | -163.50 | C3' | -155.36 | *anti* |
| 18 | C2' | *anti* | 151.02 | C3' | *anti* | -158.26 | C3' | -142.55 | *anti* |

| | | | |
|---|---|---|---|
| 19 | C3' | *anti* | -138.74 |

| Position | Nucleotide parameters for guide miR | | | | | | | | |
|---|---|---|---|---|---|---|---|---|---|
| | 4Z4D (1.60 Å) [16] | | | 6N4O (2.90 Å) [17] | | | 6NIT (3.80 Å) [18] | | |
| | Sugar pucker | Glycosidic bond | χ (°) | Sugar pucker | Glycosidic bond | χ (°) | Sugar pucker | Glycosidic bond | χ (°) |
| 1 | C2' | high-*anti* | -128.66 | C2' | high-*anti* | -117.49 | C2' | -123.66 | high-*anti* |
| 2 | **C3'** | ***anti*** | **-163.00** | **C3'** | ***anti*** | **-165.78** | C3' | **-167.28** | ***anti*** |
| 3 | **C3'** | ***anti*** | **-159.42** | **C3'** | ***anti*** | **-166.09** | C3' | **-162.40** | ***anti*** |
| 4 | **C3'** | ***anti*** | **-155.29** | **C3'** | ***anti*** | **-164.42** | C3' | **-158.22** | ***anti*** |
| 5 | **C3'** | ***anti*** | **-153.43** | **C3'** | ***anti*** | **-169.13** | C3' | **-171.20** | ***anti*** |
| 6 | **C3'** | ***anti*** | **-160.96** | **C2'** | ***anti*** | **-118.54** | C2' | **-121.99** | ***anti*** |
| 7 | **C3'** | ***anti*** | **-157.94** | **C3'** | ***anti*** | **-167.43** | C3' | **-170.51** | ***anti*** |
| 8 | **C3'** | ***anti*** | **-154.78** | **C3'** | ***anti*** | **-164.79** | C3' | **-165.32** | ***anti*** |
| 9 | **C3'** | ***anti*** | **-152.41** | **C2'** | ***syn*** | **25.41** | C3' | -143.34 | *anti* |
| 10 | C2' | high-*anti* | -102.58 | C3' | *anti* | -159.62 | **C3'** | **-151.84** | ***anti*** |
| 11 | C3' | *anti* | -165.38 | C3' | *anti* | -136.04 | **C3'** | **-163.05** | ***anti*** |
| 12 | C3' | *anti* | -166.51 | **C3'** | ***anti*** | **-144.34** | **C3'** | **-156.51** | ***anti*** |
| 13 | C3' | *anti* | -165.37 | **C3'** | ***anti*** | **-158.61** | **C3'** | **-178.03** | ***anti*** |
| 14 | C3' | *anti* | -158.94 | **C3'** | ***anti*** | **-150.41** | **C3'** | **-168.18** | ***anti*** |
| 15 | C3' | *anti* | -153.26 | C2' | *anti* | -152.75 | **C3'** | **-165.13** | ***anti*** |
| 16 | C2' | high-*anti* | -121.76 | C2' | *anti* | -156.54 | **C3'** | **-165.98** | ***anti*** |
| 17 | C3' | *anti* | -151.53 | C2' | high-*anti* | -116.59 | C3' | -161.04 | *anti* |
| 18 | C3' | -- | | C2' | *anti* | -119.01 | | | |
| 19 | C2' | high-*anti* | -67.56 | C3' | *anti* | -134.72 | | | |
| 20 | C3' | *anti* | -150.89 | | | | | | |

| Position | Nucleotide parameters for guide miR | | | | | | | | |
|---|---|---|---|---|---|---|---|---|---|
| | 6MFR (3.60 Å) [18] | | | 6MDZ (3.40 Å) [18] | | | 6MFN (2.50 Å) [18] | | |
| | Sugar pucker | Glycosidic bond | χ (°) | Sugar pucker | Glycosidic bond | χ (°) | Sugar pucker | Glycosidic bond | χ (°) |
| 1 | C2' | high-*anti* | -123.25 | C2' | high-*anti* | -123.09 | C2' | -114.54 | high-*anti* |
| 2 | **C3'** | ***anti*** | **-166.93** | **C3'** | ***anti*** | **-166.54** | **C3'** | **-163.87** | ***anti*** |
| 3 | **C3'** | ***anti*** | **-161.60** | **C3'** | ***anti*** | **-161.80** | **C3'** | **-159.89** | ***anti*** |
| 4 | **C3'** | ***anti*** | **-156.97** | **C3'** | ***anti*** | **-156.68** | **C3'** | **-157.50** | ***anti*** |
| 5 | **C3'** | ***anti*** | **-170.26** | **C3'** | ***anti*** | **-170.24** | **C3'** | **-161.38** | ***anti*** |
| 6 | **C2'** | **high-*anti*** | **-120.13** | **C2'** | **high-*anti*** | **-119.96** | **C3'** | **-158.62** | ***anti*** |
| 7 | **C3'** | ***anti*** | **-169.56** | **C3'** | ***anti*** | **-169.61** | **C3'** | **-176.50** | ***anti*** |
| 8 | **C3'** | ***anti*** | **-163.42** | **C3'** | ***anti*** | **-163.62** | C3' | -159.69 | ***anti*** |
| 9 | C2'-exo | high-*anti* | -117.97 | **C2'** | **high-*anti*** | **-120.71** | | | |
| 10 | C3' | *anti* | -145.58 | C3' | *anti* | -145.74 | | | |
| 11 | C2' | *syn* | 64.53 | C2' | *syn* | 64.41 | | | |
| 12 | **C3'** | ***anti*** | **-142.83** | **C3'** | ***anti*** | **-142.37** | | | |
| 13 | **C3'** | ***anti*** | **-154.93** | **C3'** | ***anti*** | **-154.92** | | | |
| 14 | **C3'** | ***anti*** | **-161.64** | **C3'** | ***anti*** | **-161.66** | | | |
| 15 | **C3'** | ***anti*** | **-158.26** | **C3'** | ***anti*** | **-158.24** | | | |
| 16 | **C3'** | ***anti*** | **-176.87** | **C3'** | ***anti*** | **-176.45** | | | |
| 17 | **C3'** | ***anti*** | **-165.68** | **C3'** | ***anti*** | **-165.66** | | | |
| 18 | **C3'** | ***anti*** | **-166.26** | **C3'** | ***anti*** | **-166.17** | | | |
| 19 | **C3'** | ***anti*** | **-164.08** | **C3'** | ***anti*** | **-164.02** | | | |
| 20 | C3' | *anti* | -161.34 | C3' | *anti* | -161.36 | | | |

| Position | Nucleotide parameters | | |
|---|---|---|---|
| | 7KI3 (3.00 Å) [19] | | |
| | Sugar pucker | Glycosidic bond | χ (°) |
| 1 | C2' | high-*anti* | -120.63 |
| 2 | **C3'** | ***anti*** | **-163.22** |
| 3 | **C3'** | ***anti*** | **-164.70** |
| 4 | **C3'** | ***anti*** | **-164.71** |
| 5 | **C3'** | ***anti*** | **-173.17** |
| 6 | **C2'** | **high-*anti*** | **-129.65** |
| 7 | **C3'** | ***anti*** | **-167.06** |
| 8 | **C3'** | ***anti*** | **-161.78** |
| 9 | C3' | *anti* | -163.70 |
| 10 | C3' | *anti* | -138.60 |
| 11 | C2' | high-*anti* | -123.58 |
| 12 | C3' | *anti* | -99.05 |
| 13 | C2' | high-*anti* | -126.20 |
| 14 | **C3'** | ***anti*** | **-170.64** |
| 15 | **C3'** | ***anti*** | **-168.60** |
| 16 | **C3'** | ***anti*** | **-169.75** |
| 17 | C3' | *anti* | -165.06 |
| 18 | C3' | *anti* | 178.75 |
| 19 | C3' | *anti* | -164.87 |

**Table S2**: The sequence of the miR-34a and SIRT1 mRNA, DNA template for SIRT1 mRNA and the T7 promoter.

| Sample | Sequence | nt |
|---|---|---|
| miR-34a | 5'- pU**GG**CA**G**U**G**UCUUA**G**CU**GG**UU**G**U -3' | 22 |
| SIRT1 | 5'- pGAGCUAGGACCAUUACUGCCA -3' | 21 |
| SIRT1 DNA template | 5'- *T*GGCAGTAATGGTCCTAGCTC -3' | 21 |
| T7 promoter | 5'- TATAGTGAGTCGTATTAA -3' | 18 |

* 2' O Methyl modified nucleotide, **bold $^{13}C/^{15}N$ labeled**

**Table S3**: Composition of the IVT reaction to produce SIRT1 and G$^{lab}$-miR-34a RNA.

| | | SIRT1 | miR-34a |
|---|---|---|---|
| | Stock Conc. | Volume (µL) | |
| Water | | 2800 | 400 |
| Tris-glutamate pH 8 | 500 mM | 2000 | 2000 |
| $Mg(OAc)_2$ | 250 mM | 600 | 1000 |
| DTT | 1000 mM | 100 | 100 |
| DMSO | 100% | 1000 | - |
| Spermidine | 250 mM | 1000 | 1000 |
| GMP | 100 mM | 500 | 500 |
| ATP | 100 mM | 300 | 300 |
| GTP | 100 mM | 300 | 300* |
| CTP | 100 mM | 300 | 300 |
| UTP | 100 mM | 300 | 300 |
| DNA template + T7 promoter | 20 µM | 200 | - |
| Plasmid | 20 ng/µL | - | 1000 |
| Chimera | 100 µM | - | 2000 |
| RNaseH | 1.25 mg/mL | - | 200 |
| T7 polymerase (S43Y) | 3.7 mg/mL | 600 | 600 |
| | Total | 10,000 | 10,000 |

* $^{13}C/^{15}N$ labeled

**Table S4**: Concentrations of hAgo2 protein and the amount of $G^{lab}$-miR-34a and SIRT1 used for the DNP MAS NMR experiments.

| **Sample** | **Amount** |
|---|---|
| miR-34a | 80 nmol |
| miR-34a:SIRT1 | 80 nmol |
| hAgo2* (total amount)<br>miR-34a‡ : hAgo2 | 3 nmol<br>1.45 nmol : 1.5 nmol |
| miR-34a‡ : hAgo2 : SIRT1 | 1.45 nmol : 1.5 nmol : 1.5 nmol |

* Quantification from combined Bradford assay and SDS PAGE,

‡ From Northern blot

**Table S5**. Important experimental parameters for the DNP MAS NMR for different samples studied.

| Experiment | Sample | SW (ppm) (F1/F2) | TD1 | AQ (ms) | NS | D1 (s) |
|---|---|---|---|---|---|---|
| $^{13}C$ - $^{13}C$ DARR | $G^{lab}$-miR-34a | 300/300 | 214 | 21.25 | 8 | 4 |
| | $G^{lab}$-miR-34a:SIRT1 | 300/300 | 196 | 21.25 | 8 | 4 |
| | hAGO2:$G^{lab}$-miR-34a | 300/300 | 214 | 21.25 | 256 | 4 |
| | hAGO2:$G^{lab}$-miR-34a:SIRT1 | 300/300 | 196 | 34.00 | 128 | 4 |
| $^{13}C$ - $^{31}P$ TEDOR | $G^{lab}$-miR-34a | 300/37.04 | 96 | 21.25 | 8 | 3 |
| | $G^{lab}$-miR-34a:SIRT1 | 300/24.68 | 32 | 21.25 | 16 | 3 |
| | hAGO2:$G^{lab}$-miR-34a | 300/37.04 | 128 | 21.25 | 220 | 3.5 |
| | hAGO2:$G^{lab}$-miR-34a:SIRT1 | 300/37.04 | 128 | 21.25 | 160 | 3 |

**Table S6**. Gaussian fit parameters of the positive projection shown in Figure S9a for hairpin $G^{lab}$-miR-34a. The C-C correlation associated with the fit is denoted. The fit for C8 – C1’ positive projection for C3’-endo was fitted with two gaussians. The 1σ standard deviation is shown for the amplitude, δ and σ.

| C3'-endo | Amplitude | Amplitude Std.Dev | δ (ppm) | δ (ppm) Std.Dev | σ | σ Std.Dev |
|---|---|---|---|---|---|---|
| C5 – C1’ | 150377548 | 717109 | 118.51 | 0.01 | 1.24 | 0.01 |
| C8 – C1’ | 104515868 | 6842751 | 136.19 | 0.04 | 1.17 | 0.02 |
| C8 – C1’ | 104885628 | 6934953 | 138.77 | 0.11 | 1.60 | 0.00 |
| C2 – C1’ | 202296129 | 2053961 | 152.19 | 0.01 | 1.39 | 0.01 |
| C4 – C1’ | 174271258 | 2180468 | 156.00 | 0.07 | 1.45 | 0.02 |
| C6 – C1’ | 124614703 | 785553.4 | 161.18 | 0.01 | 1.17 | 0.01 |
| C2'-endo | | | | | | |
| C5 – C1’ | 32727456 | 779915.8 | 118.84 | 0.05 | 1.80 | 0.02 |
| C8 – C1’ | 32706635 | 779913.1 | 141.83 | 0.05 | 1.80 | 0.11 |
| C2/4 – C1’ | 49496716 | 633876.5 | 155.00 | 0.02 | 1.19 | 0.02 |
| C6 – C1’ | 25211575 | 614586.8 | 161.54 | 0.03 | 1.12 | 0.03 |

δ = Center of the gaussian fit or the chemical shift of the resonance,

σ = Width of the gaussian fit, where the full width at half maxima can be calculated with $2\sigma\sqrt{2\ \ln 2}$

**Table S7**. Gaussian fit parameters of the positive projection shown in Figure S9b for duplex $G^{lab}$-miR-34a:SIRT1. The C-C correlation associated with the fit is denoted. The fit for C8 – C1’ positive projection for C2’- and C3’-endo was fitted with two gaussians. The 1σ standard deviation is shown for the amplitude, δ and σ.

| C3'-endo | Amplitude | Amplitude Std.Dev | δ (ppm) | δ (ppm) Std.Dev | σ | σ Std.Dev |
|---|---|---|---|---|---|---|
| C5 – C1’ | 1207972036 | 1754538 | 118.69 | 0.00 | 1.29 | 0.00 |
| C8 – C1’ | 1168919420 | 71965584 | 136.15 | 0.02 | 1.70 | 0.02 |
| C8 – C1’ | 540859970 | 72916030 | 138.92 | 0.33 | 2.50 | 0.00 |
| C2 – C1’ | 1677936747 | 2636214 | 152.02 | 0.00 | 1.52 | 0.00 |
| C4 – C1’ | 1068962518 | 2659186 | 156.66 | 0.00 | 1.31 | 0.00 |
| C6 – C1’ | 930657012 | 1807858 | 161.71 | 0.00 | 1.18 | 0.00 |
| C2'-endo | | | | | | |
| C5 – C1’ | 143884193 | 1204569 | 118.25 | 0.01 | 1.43 | 0.01 |
| C8 – C1’ | 151013082 | 3292161 | 139.2 | 0.25 | 1.80 | 0.19 |
| C8 – C1’ | 70698581 | 3104658 | 143.08 | 0.05 | 1.23 | 0.04 |
| C2 – C1’ | 54240711 | 17411494 | 151.8 | 0.17 | 1.80 | 0.12 |
| C4 – C1’ | 287878777 | 17492779 | 155.12 | 0.09 | 1.71 | 0.04 |
| C6 – C1’ | 135315984 | 1189680 | 161.42 | 0.01 | 1.20 | 0.01 |

δ = Center of the gaussian fit or the chemical shift of the resonance,
σ = Width of the gaussian fit, where the full width at half maxima can be calculated with $2\sigma\sqrt{2\ \ln 2}$

**Table S8**. Gaussian fit parameters of the positive projection shown in Figure S9c for hAgo:$G^{lab}$-miR-34a binary complex. The C-C correlation associated with the fit is denoted. The fit for C8 – C1' and C5 – C1' positive projection for C2'-endo was fitted with two gaussians. The 1σ standard deviation is shown for the amplitude, δ and σ.

| C3'-endo | Amplitude | Amplitude Std.Dev | δ (ppm) | δ (ppm) Std.Dev | σ | σ Std.Dev |
|---|---|---|---|---|---|---|
| C5 – C1' | 581171077 | 5064129 | 118.70 | 0.01 | 1.50 | 0.01 |
| C8 – C1' | 812807277 | 5625291 | 136.69 | 0.01 | 1.86 | 0.01 |
| C2 – C1' | 993072894 | 8327889 | 152.68 | 0.02 | 1.87 | 0.02 |
| C4 – C1' | 493379596 | 6798432 | 156.71 | 0.01 | 0.90 | 0.01 |
| C6 – C1' | 596528350 | 4827396 | 162.32 | 0.01 | 1.36 | 0.01 |
| C2'-endo | | | | | | |
| C5 – C1' | 105510461 | 22664718 | 117.90 | 0.06 | 0.86 | 0.08 |
| C5 – C1' | 183918812 | 23106302 | 118.78 | 0.17 | 1.80 | 0.06 |
| C8 – C1' | 303771583 | 22981619 | 138.70 | 0.07 | 1.31 | 0.05 |
| C8 – C1' | 176457584 | 23642103 | 142.16 | 0.27 | 1.80 | 0.06 |
| C2 – C1' | 421242581 | 47471088 | 154.84 | 0.19 | 1.52 | 0.10 |
| C4 – C1' | 256741260 | 46592682 | 157.00 | 0.03 | 0.97 | 0.05 |
| C6 – C1' | 374199813 | 5937404 | 162.26 | 0.03 | 1.80 | 0.02 |

δ = Center of the gaussian fit or the chemical shift of the resonance,
σ = Width of the gaussian fit, where the full width at half maxima can be calculated with $2\sigma\sqrt{2\ \ln 2}$

**Table S9**. Gaussian fit parameters of the positive projection shown in Figure S9d for hAgo:G$^{lab}$-miR-34a:SIRT1 ternary complex. The C-C correlation associated with the fit is denoted. The fit for C8 – C1’ positive projection for C3’-endo was fitted with two gaussians. The 1σ standard deviation is shown for the amplitude, δ and σ.

| C3'-endo | Amplitude | Amplitude Std.Dev | δ (ppm) | δ (ppm) Std.Dev | σ | σ Std.Dev |
|---|---|---|---|---|---|---|
| C5 – C1’ | 306598070.26 | 4125455 | 118.76 | 0.02 | 1.54 | 0.02 |
| C8 – C1’ | 295812312.18 | 4719104 | 137.00 | 0.57 | 2.00 | 90.15 |
| C8 – C1’ | 318418967.06 | 12611186 | 153.00 | 0.08 | 1.54 | 0.06 |
| C2 – C1’ | 88881891.05 | 19156041 | 155.41 | 0.07 | 0.70 | 0.07 |
| C4 – C1’ | 230734977.26 | 14333016 | 157.40 | 0.04 | 1.08 | 0.06 |
| C6 – C1’ | 215609163.03 | 4649050 | 163.00 | 0.04 | 1.67 | 0.04 |
| C2'-endo | | | | | | |
| C5 – C1’ | 244285316.68 | 4251361 | 118.00 | 0.03 | 1.83 | 0.04 |
| C8 – C1’ | 326825235.43 | 4438360 | 140.66 | 0.03 | 2.00 | 0.03 |
| C2/4 – C1’ | 458897767.71 | 4159219 | 156.50 | 0.01 | 1.74 | 0.02 |
| C6 – C1’ | 104696571.41 | 2763394 | 162.92 | 0.02 | 0.77 | 0.02 |

δ = Center of the gaussian fit or the chemical shift of the resonance,

σ = Width of the gaussian fit, where the full width at half maxima can be calculated with $2\sigma\sqrt{2\ \ln 2}$

**Table S10**. Gaussian fit parameters for individual peaks in the C8 to C1' correlation from binary complex as shown in Figure 1g of the main text. The color coding indicates the resonances having a distribution of both C2'- and C3'-endo conformation where each color belongs to one guanosine. See the main text for details. The percentage distribution for these resonances is also denoted.

| Percentage distribution of puckering per guanosine | Guanosine | Amplitude | Amplitude Std.Dev | δ (ppm) | δ (ppm) Std.Dev | σ | σ Std.Dev |
|---|---|---|---|---|---|---|---|
| | | **C3'-endo** | | | | | |
| | $G^{C3'}$-1 | 35362286.25 | 4487571.37 | 132.99 | 0.024 | 0.56 | 0.03 |
| | $G^{C3'}$-2 | 14882772.06 | 990655.71 | 133.95 | 0.006 | 0.21 | 0.01 |
| | $G^{C3'}$-3 | 446618474.1 | 67333035.67 | 135.95 | 0.194 | 1.33 | 0.13 |
| | $G^{C3'}$-4 | 101962585.4 | 37201905.02 | 136.83 | 0.047 | 0.52 | 0.05 |
| 36 ± 0.04% | $G^{C3'}$-5 | 68722567.42 | 9648430.13 | 137.79 | 0.019 | 0.37 | 0.03 |
| | $G^{C3'}$-6 | 35881273.54 | 8889036.09 | 138.44 | 0.025 | 0.29 | 0.02 |
| 68 ± 0.06% | $G^{C3'}$-7 | 87645774.79 | 16813029.97 | 139.15 | 0.085 | 0.72 | 0.05 |
| 78 ± 0.01% | $G^{C3'}$-8 | 45233526.91 | 2044210.78 | 141.49 | 0.029 | 0.8 | 0.12 |
| 41 ± 0.01% | $G^{C3'}$-9 | 34561192.66 | 797799.70 | 143.62 | 0.009 | 0.48 | 0.01 |
| | | **C2'-endo** | | | | | |
| 64 ± 0.04% | $G^{C2'}$-1 | 41077868.69 | 6736994.79 | 137.19 | 0.04 | 0.43 | 0.02 |
| | $G^{C2'}$-2 | 144605238.9 | 14126248.26 | 138.35 | 0.01 | 0.61 | 0.05 |
| 32 ± 0.06% | $G^{C2'}$-3 | 41932663.01 | 8827100.37 | 139.25 | 0.02 | 0.3 | 0.02 |
| | $G^{C2'}$-4 | 20154794.25 | 3327654.41 | 139.77 | 0.02 | 0.23 | 0.01 |
| | $G^{C2'}$-5 | 113812454.3 | 2765198.55 | 140.64 | 0.01 | 0.65 | 0.02 |
| 22 ± 0.04% | $G^{C2'}$-6 | 13056750.78 | 3321645.81 | 141.85 | 0.05 | 0.24 | 0.02 |
| | $G^{C2'}$-7 | 28429262.94 | 3174327.76 | 142.35 | 0.03 | 0.29 | 0.02 |
| 59 ± 0.01% | $G^{C2'}$-8 | 50540560.02 | 683349.13 | 143.54 | 0.01 | 0.6 | 0.01 |

**Table S11**. Gaussian fit parameters for individual peaks in the C8 to C1' correlation from ternary complex as shown in Figure 1g of the main text. The color coding indicates the resonances having a distribution of both C2'- and C3'-endo conformation where each color belongs to one guanosine. The percentage distribution for these resonances is also denoted.

| Percentage distribution of puckering per guanosine | Guanosine | Amplitude | Amplitude Std.Dev | δ (ppm) | δ (ppm) Std.Dev | σ | σ Std.Dev |
|---|---|---|---|---|---|---|---|
| | | **C3'-endo** | | | | | |
| | $G^{C3'}$-1 | 13118849.35 | 2057899.84 | 134.96 | 0.02 | 0.28 | 0.02 |
| | $G^{C3'}$-2 | 52998846.83 | 4907985.53 | 135.85 | 0.016 | 0.44 | 0.03 |
| | $G^{C3'}$-3 | 1226420.79 | 364570.71 | 136.55 | 0.016 | 0.1 | 0.02 |
| | $G^{C3'}$-4 | 114547204.9 | 5402819.91 | 137.12 | 0.014 | 0.6 | 0.03 |
| 25 ± 0.05% | $G^{C3'}$-5 | 10275594.06 | 2792725.8 | 138.51 | 0.009 | 0.24 | 0.02 |
| 76 ± 0.03% | $G^{C3'}$-6 | 44689089.43 | 5590817.52 | 139.09 | 0.063 | 0.59 | 0.05 |
| 55 ± 0.06% | $G^{C3'}$-7 | 47116731.12 | 1371454.08 | 141.01 | 0.015 | 0.64 | 0.02 |
| | | **C2'-endo** | | | | | |
| 75 ± 0.05% | $G^{C2'}$-1 | 30610504.4 | 759819.13 | 138.48 | 0.006 | 0.35 | 0.01 |
| 24 ± 0.03% | $G^{C2'}$-2 | 14058042.8 | 1246870.33 | 139.22 | 0.005 | 0.21 | 0.01 |
| | $G^{C2'}$-3 | 114852011 | 6017730.44 | 140.14 | 0.018 | 0.62 | 0.03 |
| 45 ± 0.06% | $G^{C2'}$-4 | 37996674.4 | 9999346.34 | 141.26 | 0.043 | 0.37 | 0.05 |
| | $G^{C2'}$-5 | 10767374.5 | 4104531.27 | 141.68 | 0.01 | 0.2 | 0.02 |
| | $G^{C2'}$-6 | 58062495.5 | 3143435.1 | 142.46 | 0.02 | 0.57 | 0.03 |
| | $G^{C2'}$-7 | 18263629.5 | 966276.56 | 143.63 | 0.007 | 0.3 | 0.01 |

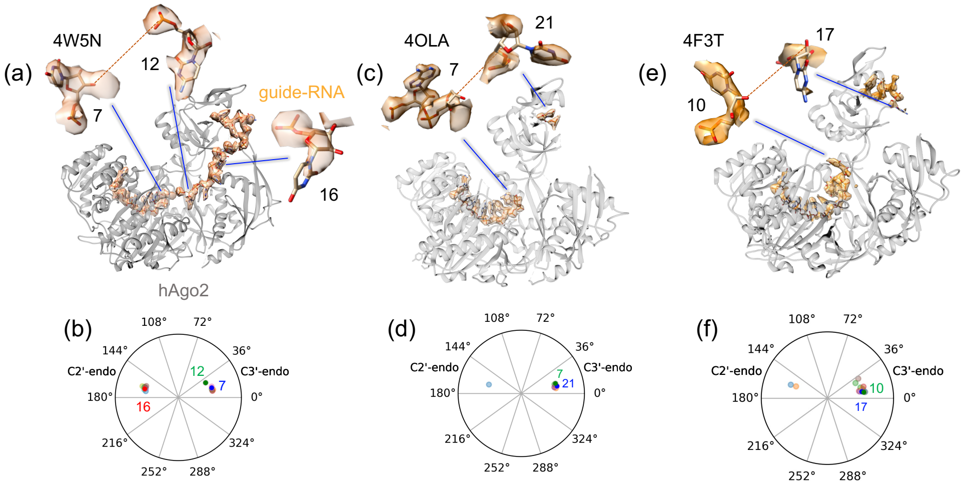

**Figure S1.** miR conformation within hAgo2 from crystal structures 4W5N[6] (a), 4OLA[5] (c) and 4F3T[15] (e) with protein in grey and RNA in orange. The $2F_0 - F_c$ electron density at 1σ of miR is shown in light orange. The dashed red line represents missing segments. Pseudo-rotation cycle corresponding to each crystal structure (panels b, d, and f) shows the distribution of ribose C2'- and C3'-endo pucker. Nucleotides showing the challenge of modelling ribose puckering and nucleobase orientation due to poorly defined electron density is shown above each crystal structures and numbered in the pseudo-rotation cycles.

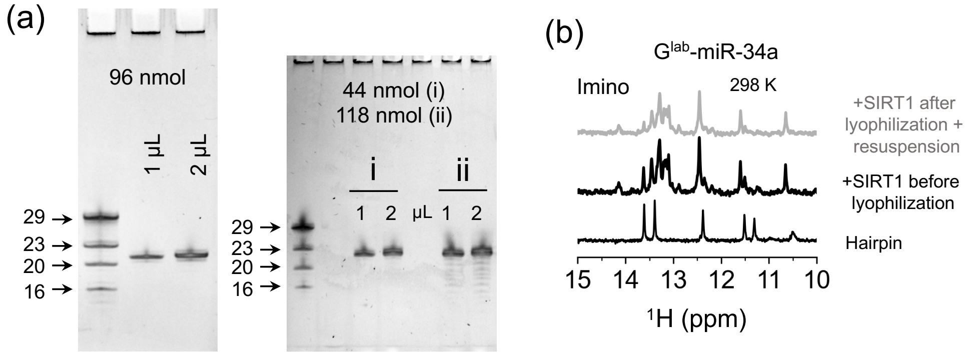

**Figure S2**. **RNA production and purification.** (a) Quality check of purified RNA on 20% denaturing polyacrylamide gel. The left panel shows the purity of 1 and 2 μL of 21 nt SIRT1 RNA from a solution of 9.6 pmol/μL. The total yield in nmols is shown on the gel. The right panel shows the purity of $G^{lab}$-miR-34a after two IVT reactions (i and ii). 1 and 2 μL from a solution of 4.4 pmol/μL and 11.8 pmol/μL was loaded for each reaction respectively and is shown on the gel. The total yield for each reaction were 44 and 118 nmols respectively. The ladder with RNA sizes of 29, 23, 20 and 16 nucleotides are also shown for both the panels. (b) Solution state 1D $^{1}H$ spectra at 298 K of $G^{lab}$-miR-34a:SIRT1 complex before (black) and after lyophilization + resuspension (grey) depicting that the overall spectrum do not change due to lyophilization. The spectrum of the hairpin after lyophilization and resuspension is similar to the previous report[7] thereby, suggesting that its structure is also not affected by this treatment.

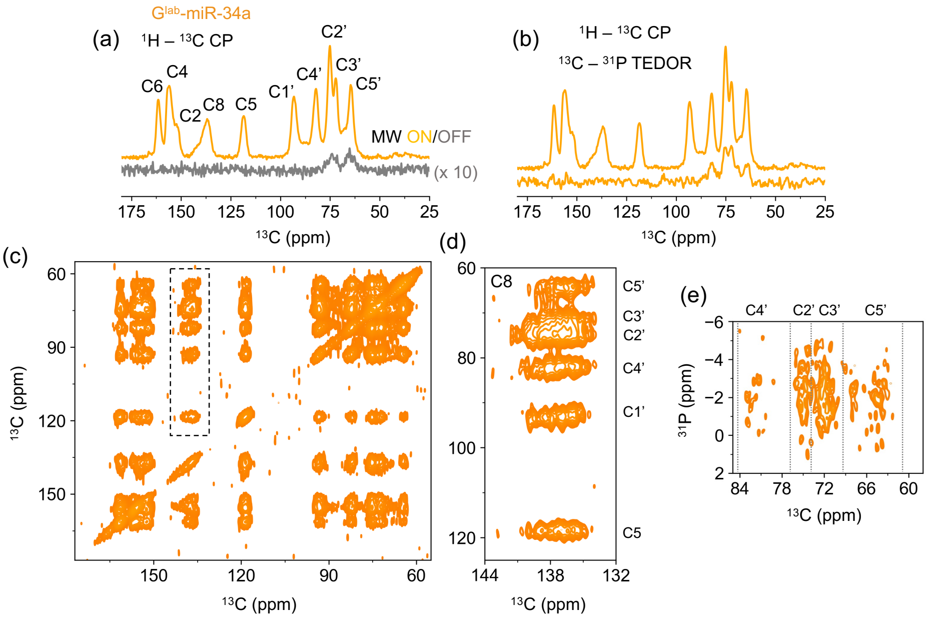

**Figure S3**. **1D and 2D spectra of G$^{lab}$-miR-34a hairpin.** (a) 1D $^{1}$H-$^{13}$C CP spectra, exhibiting the ribose- and nucleobase carbons (C1'-C5' and C2, C4, C5, C6, C8, structure Figure 1c) from guanosines. Microwave off spectrum is shown in grey with a 10x increased intensity showing the residual glycerol signal (b) Overlay of 1D $^{1}$H-$^{13}$C CP (top) and (bottom) 1D $^{31}$P-$^{13}$C TEDOR showing that the ribose carbons C2', C3', C4', and C5' near the phosphorus can be selectively observed. (c) 2D $^{13}$C-$^{13}$C DARR spectra, with a zoomed-in (dashed box) area showing the correlation from C8 to C5 and all ribose carbons in panel (d). 8 distinguishable guanosine signals from the C8 to C5 correlation are marked with solid lines, presenting the 8 labeled gunaosines. (e) 2D $^{31}$P-$^{13}$C TEDOR spectrum, revealing a $^{31}$P chemical shift dispersion of approximately 6 ppm. Different regions for C5', C4', C3' and C2' are marked with dashed grey line.

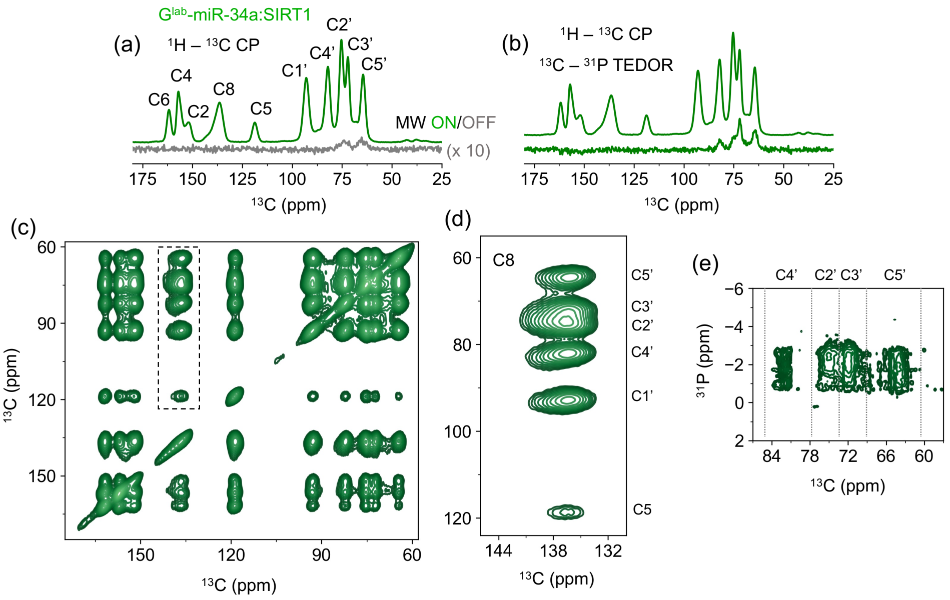

**Figure S4. 1D and 2D spectra of G$^{lab}$-miR-34a:SIRT1 duplex** (a) 1D $^{1}$H-$^{13}$C CP spectra that exhibit the ribose (C1'-C5') and nucleobase (C2, C4, C5, C6, C8) carbons of guanosine. Microwave off spectrum is shown in grey with a 10x increased intensity showing the residual glycerol signal. (b) Comparison of $^{1}$H-$^{13}$C CP and $^{13}$C-$^{31}$P TEDOR experiment showing that the ribose carbons except C1' can be selectively observed similar. (c) 2D $^{13}$C-$^{13}$C DARR spectra, with a zoomed-in (dashed box) area showing the correlation from C8 to C5 and all ribose carbons in panel (d). In comparison with Figure S4c, the 2D DARR spectrum of the duplex shows line broadening caused by the large chemical shift anisotropy. (e) The$^{13}$C-$^{31}$P TEDOR spectrum showing the $^{31}$P chemical shift has less dispersion (~3.5 ppm) compared to free G$^{lab}$-miR-34a due to the well-defined duplex structure. Different regions for C5', C4', C3' and C2' are marked with dashed grey line.

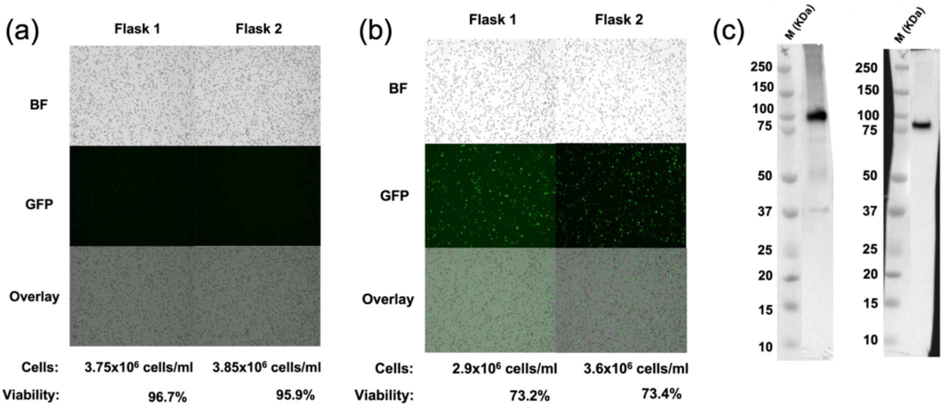

**Figure S5**. **Insect cell protein expression and baculovirus production.** Content of two 3 L Erlenmeyer flasks labeled as flask 1 and flask 2 (each containing 1 L of media) are shown here. After transfection with P2 baculovirus stock, Sf9 cells and their expression levels were monitored at 24 h (a) and 48 h (b). Microscopy snapshots including images captured under bright field (BF), GFP filter (GFP), and an overlay of BF and GFP (Overlay) are shown. Cells per mL and the viability in percentage are also reported. (c) Western blots using two different antibodies targeting either the N-domain which includes the N-terminal, Linker-1, and PAZ (right panel) or the C-domain which includes, Linker-2, MID and PIWI (left panel)[5]. Details about the antibodies can be found in the material and methods.

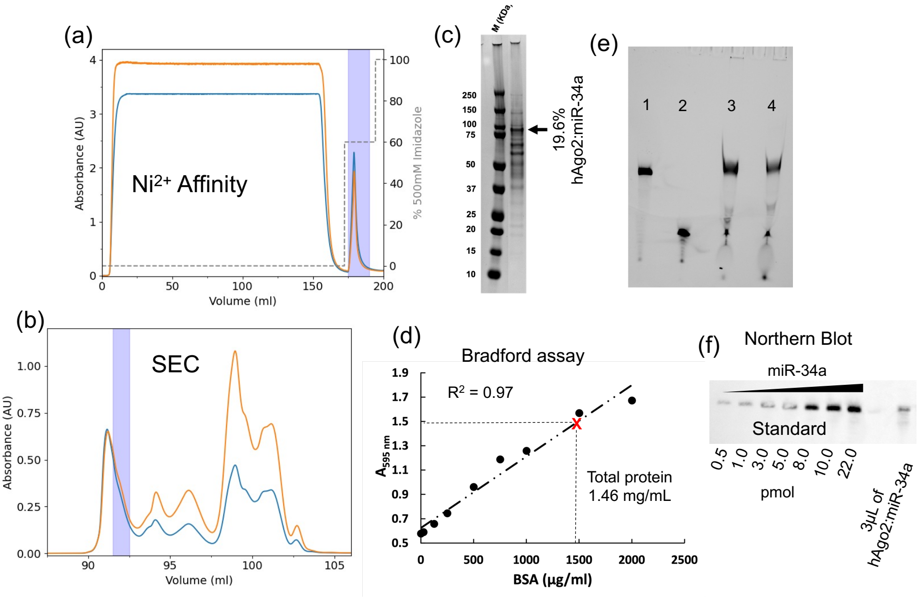

**Figure S6**. **Protein purification and quantification**. (a) Chromatogram of hAgo2 from SF9 insect cell through the HisTrap-$Ni^{2+}$ column monitored at 280 (blue) and 260 (orange) nm wavelengths. The step gradient of elution buffer with 500 mM Imidozole is shown as dashed lines. Elution was done at 300 mM imidazole. The fractions pooled for further purification is highlighted in violet (b) Size exclusion chromatogram of hAgo2 loaded with $G^{lab}$-miR-34a and after TEV protease digestion monitored at 280 (blue) and 260 (orange) nm wavelengths. The fractions collected are highlighted in violet. (c) SDS-PAGE of the hAgo2:$G^{lab}$-miR-34a complex after size exclusion chromatography (SEC). The molecular weight ladder is shown with their respective sizes in kDa. The hAgo2 is shown with an arrow which is 19.6% of the total protein. (d) Bradford assay for total protein concentration. The estimated protein absorbance at 595 nm and corresponding concentration is marked as red x on a BSA standard curve. The concentration from the linear fit of the BSA standard curve (equation 1) with $R^2$ value of 0.97 gives a total protein of 1.46 mg/mL. (e) The slicing activity assay of the hAgo2:$G^{lab}$-miR-34a complex before (lane 3) and after (lane 4) lyophilization. Lane 1 and 2 represents the control with Cy3 labeled 34nt and 15nt product respectively. It shows that the activity is not affected due to lyophilization. (f) Northern blot quantification of $G^{lab}$-miR-34a loaded in hAgo2. The amount loaded for the miR-34a standard curve and the lane with 3µL of hAgo2:miR-34a complex is denoted. The estimated amount (equation 2) of miR-34a loaded is 2.9 pmol/µL.

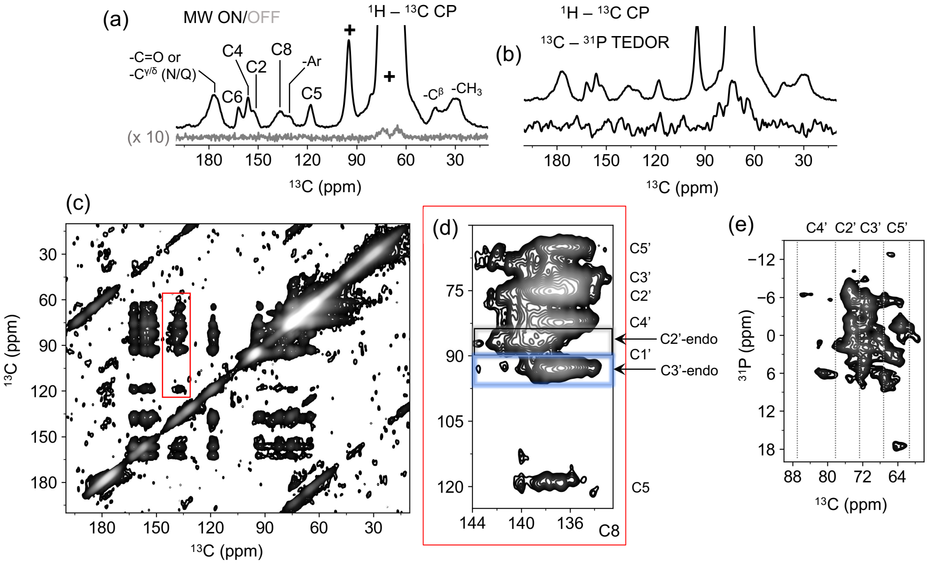

**Figure S7**. **1D and 2D spectra of hAgo2:G$^{lab}$-miR-34a binary complex (black).** (a) 1D $^{1}$H-$^{13}$C CP spectrum of binary complex. Microwave off spectrum is shown in grey with a 10x increased intensity showing the residual glycerol signal. In addition to the signals from guanosines, carbon resonances from the carbonyl and/or C$^{\gamma}$ (Asn) or C$^{\delta}$ (Gln) residues, aromatic ring carbons (Ar), aliphatic -C$^{\beta/\gamma}$, and -$CH_3$ carbon signals of the hAgo2 protein are also observed. The signals originating from cryoprotectants (trehalose and mannitol) and residual glycerol from the sample preparation, are indicated with "+". (b) 1D $^{31}$P-$^{13}$C TEDOR spectrum compared to the CP, showing that the ribose carbon can be observed even though the region is overrepresented with cryoprotectants and glycerol. (c) 2D $^{13}$C-$^{13}$C DARR spectrum of hAgo2:G$^{lab}$-miR-34a complex. The red box is zoomed in panel (d) showing the C8 correlations to C5 and all the ribose carbons. The C2'- (grey box) and C3'-endo (blue box) regions from the C8 to C1' correlation are denoted. (e) The 2D $^{31}$P-$^{13}$C TEDOR spectrum reveals that the G$^{lab}$-miR-34a's $^{31}$P resonance exhibits a broader range of approximately 16 ppm when it is bound within hAgo2, compared to around 6 ppm when it is in its unbound state (Fig. S3). Different regions corresponding to the ribose carbon C5', C3', C2' and C4' are denoted with dashed lines.

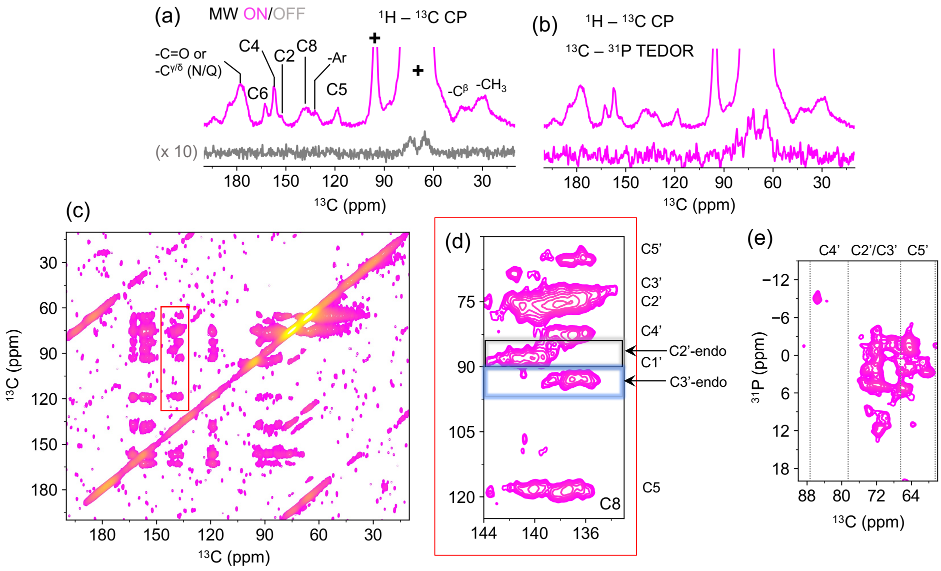

**Figure S8. 1D and 2D spectra of hAgo2:G$^{lab}$-miR-34a:SIRT1 ternary complex.** 1D $^{1}$H-$^{13}$C CP spectrum of ternary complex. Microwave off spectrum is shown in grey with a 10x increased intensity showing the residual glycerol signal. Similar to the binary complex, resonances from both protein and RNA could be identified. (b) $^{13}$C-$^{31}$P TEDOR experiment allows for the selective observation of the ribose carbons. (c) 2D $^{13}$C-$^{13}$C DARR spectrum with the red box showing the zoomed in region of C8 correlation to C5 and all the ribose in (d). The C2'- (grey box) and C3'-endo (blue box) regions from the C8 to C1' correlation are denoted. (e) The 2D $^{31}$P-$^{13}$C TEDOR where the spread of $^{31}$P resonances is similar to hAgo2:G$^{lab}$-miR-34a complex, suggesting that the duplex release phenomenon[18] is suppressed in this case. However, differences are observed in the absolute resonance position and is discussed in the main text. Different regions corresponding to the ribose carbon C5', C3', C2' and C4' are denoted with dashed lines. Contrary to the binary complex, no correlation from protein (carbonyl carbon) to RNA is observed in the ternary complex (Figure 1i in the main text). Since there is not duplex release it is expected that there will be some DARR correlation between the amino acid side chain and the ribose. However, the chemical shift of the amino acid side chain overlap with the labelled ribose and residual trehalose, mannitol, and glycerol. Additionally, the signal intensity will be very small compared to the labelled ribose hence these correlations cannot be observed unambiguously.

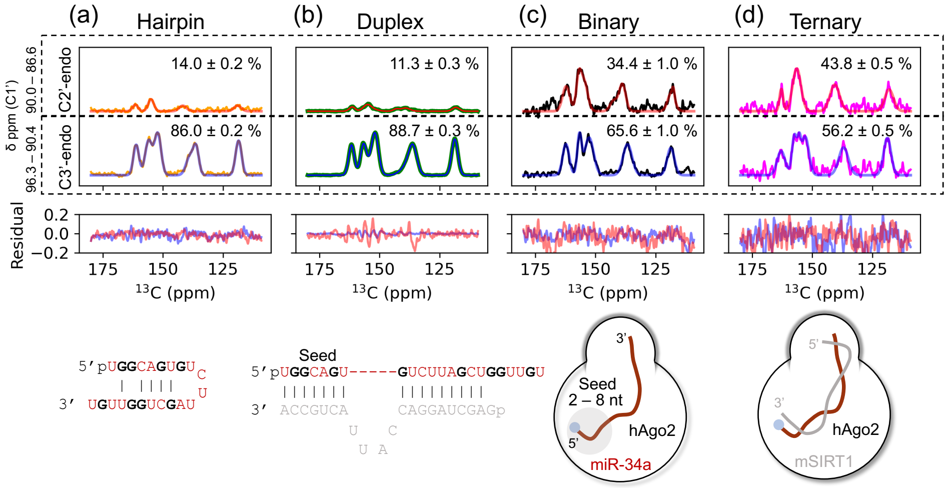

**Figure S9**. Gaussian fits of the positive projection of the C1' regions corresponding to C2'- (90.0 – 86.6 ppm) and C3'-endo (96.3 – 90.4 ppm) conformers from the DARR spectra of hairpin (orange, a), duplex (green, b), binary (black, c) and ternary (magenta, d) complex. The red and blue solid lines are the fits and the residuals are shown for each sample together, presented in Table S6 – S9. The chemical shift region used for the projection is denoted on the left side. The overall distribution in percent for each the conformers is shown on the top of the graph. The difference in noise level for different sample is due to different concentrations. The secondary structure for hairpin and duplex, and schematic representation of the binary and ternary complex are shown below each graph. The seed region is denoted in the duplex and binary complex.

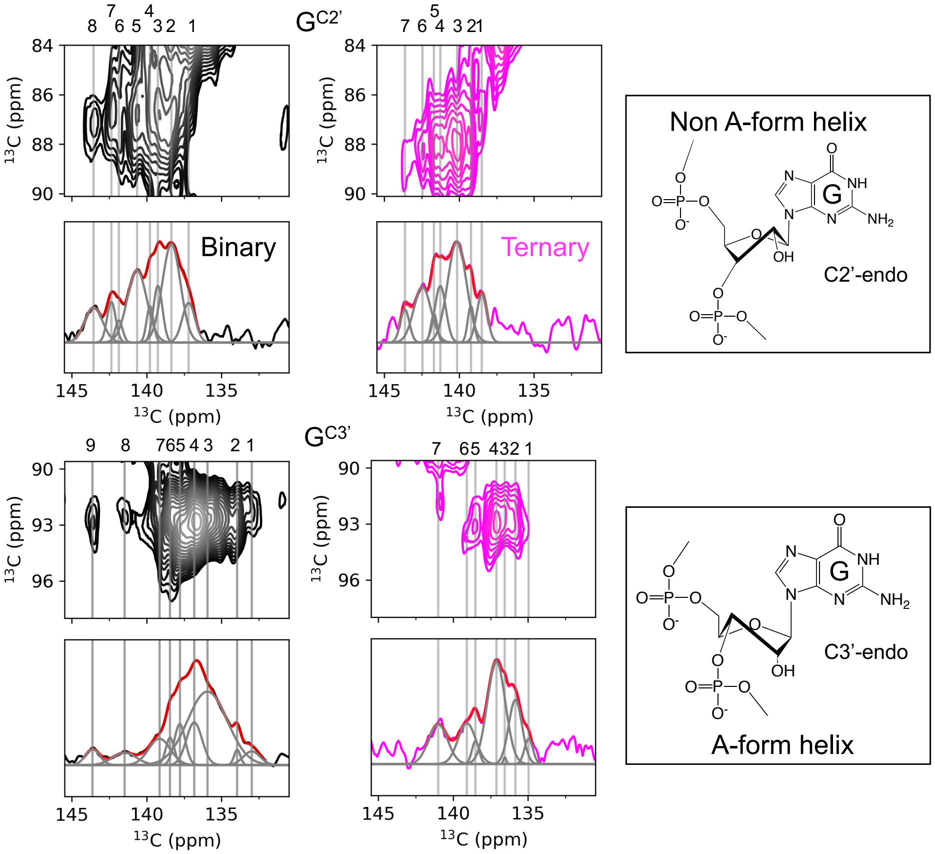

**Figure S10.** Gaussian fits of the positive projection of C1' signal in the C2' and C3'-endo conformation for binary (black) and ternary (magenta) complex. Solid grey lines show the position of the signals. The cumulative fit of the deconvoluted signals (red) is overlayed on the 2D contour (gray) to indicate the quality of the individual fits (gray) of the positive projection, details are tabulated in Table S10 – S11. The guanosines for C2'($G^{C2'}$) and C3' ($G^{C3'}$)-endo are numbered and denoted over the solid lines. Chemical structures indicating the conformers are provided on the right.

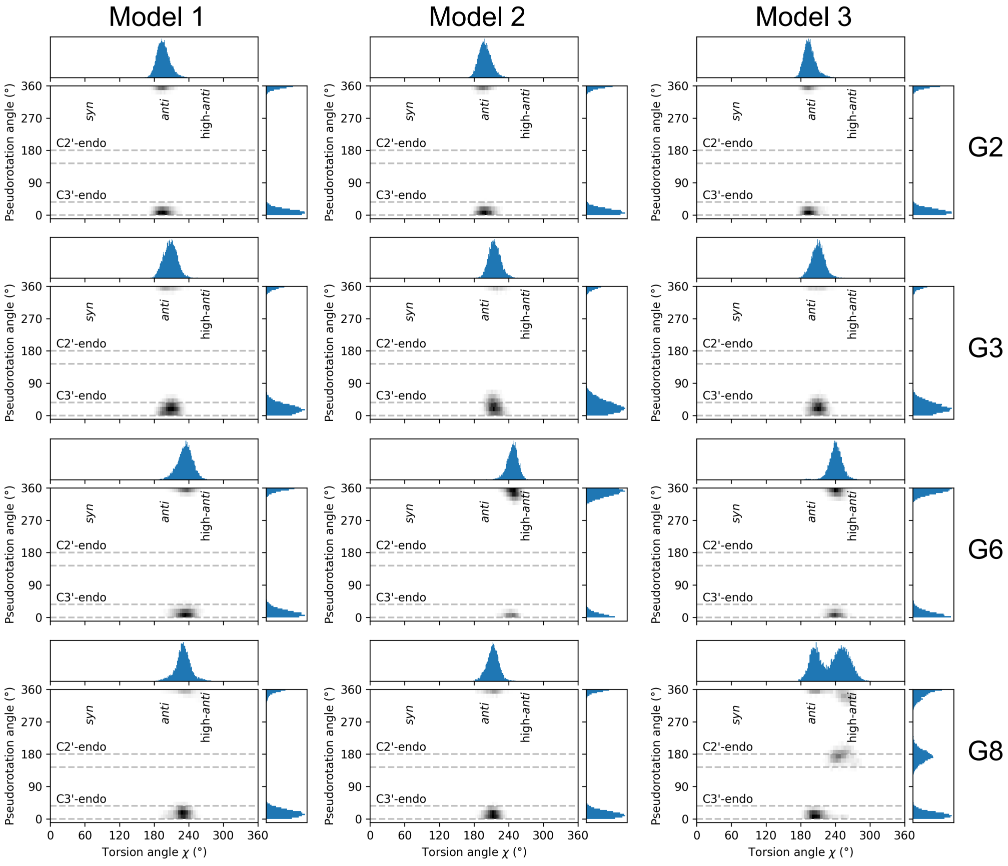
Model 1
Model 2
Model 3
G2
G3
G6
G8
Pseudorotation angle (°)
Torsion angle χ (°)
syn
anti
high-anti
C2'-endo
C3'-endo
0
60
120
180
240
300
360
0
90
180
270
360

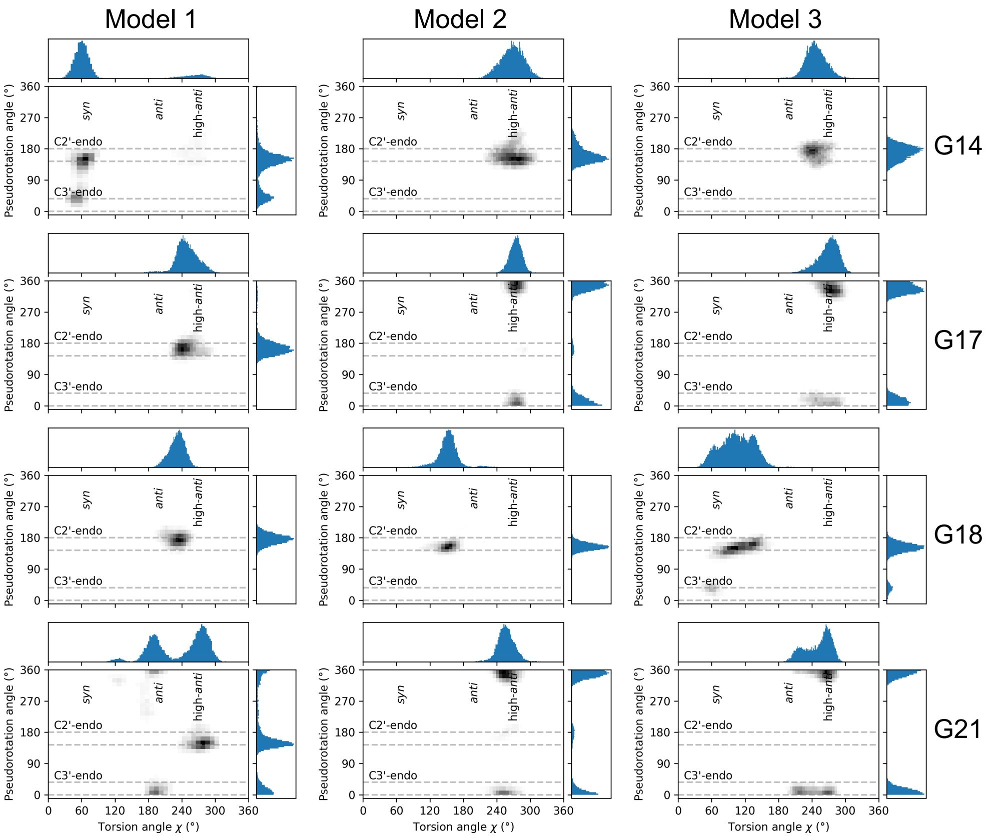

**Figure S11:** Correlation plots of the sugar pucker (pseudo rotation angle) and the glycosidic bond torsion angle (χ) for the guanosines in the binary complex from three different starting structures simulated for 400 ns each. The ribose C3'- and C2'-endo puckering area is marked with dashed lines. The regions for the syn, anti and high-anti conformation is shown. The histogram of the distribution is shown at the top and bottom for the χ angle and pseudo rotation angle respectively. The numbering of guanosines is shown at the right side. It is observed that seed G's have C3'-endo distribution, while other guanosines can attain several different conformations as a combination of the χ angle and ribose sugar puckering.

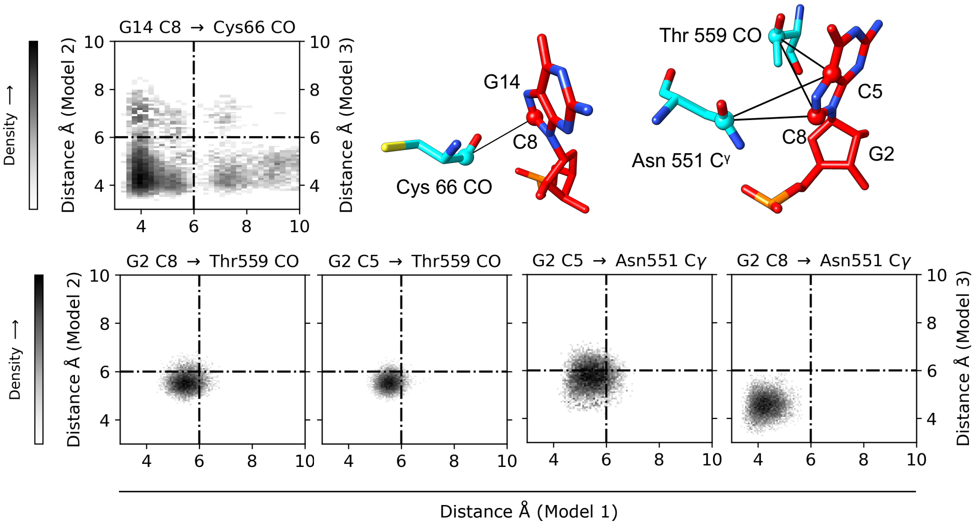

**Figure S12. Protein-RNA interactions observed in MD simulations.** Density plots of the distances between carbons of potential amino acids and C5/C8 of guanosines from MD simulations that contribute to the DARR correlations in the binary complex. Three different starting structures are named as model 1, model 2 and model 3 in their respective axes. Gradient from low (white) to high (black) density is shown on the left side of the plots. The 6 Å cut-off distance of below which a DARR correlation is observed[20], is shown as dashed-dot lines. The amino acids satisfying the 6 Å cut-off criteria from the trajectories of the three starting structures are Cys66, ASN551, and Thr559 for G14 and G2, respectively. The positions of the carbons from respective amino acids in cyan (Cys66 CO, Asn551 $C^{\gamma}$ and Thr559 CO) and guanosine in red (G2 C8 and G14 C5/C8) of miR-34a from the time-averaged model is shown (upper right). The relevant distances between the carbons are indicated with solid black lines.

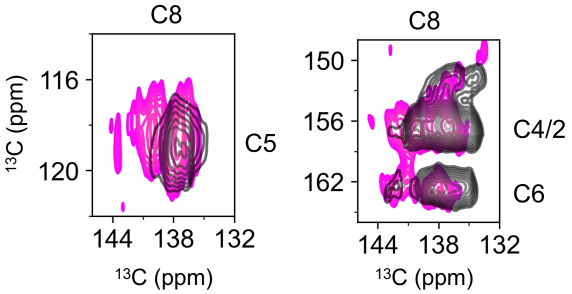

**Figure S13.** DARR regions of nucleobase correlation between C8 to C6, C4/2 (right) and C5 (left) highlighting the changes between binary (black) and ternary (magenta) complexes. The chemical shift changes suggest binding to the SIRT1 mRNA. Typically these RNA:RNA interactions involve base-pairing, leading to an increase in distance of the base previously interacting with the protein and therefore a loss of visible contacts in the DARR spectrum of the ternary complex.

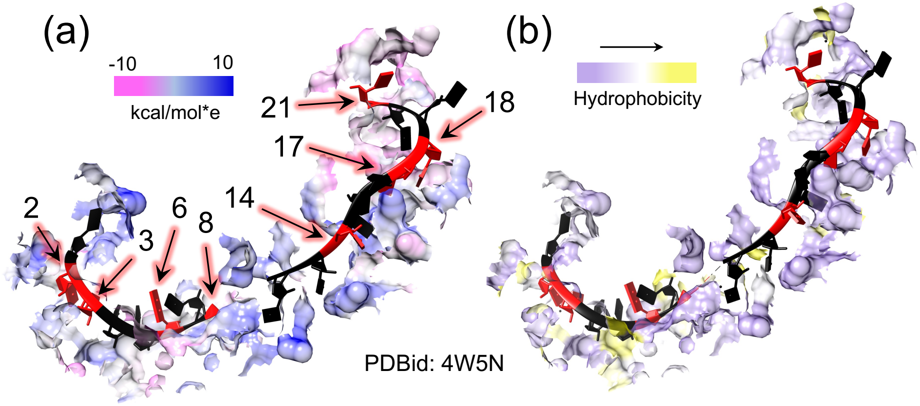

**Figure S14:** (a) Variation of electrostatic potential is shown as a surface calculated by the Adaptive Poisson–Boltzmann Solver[21] for the amino acids within 5 Å of the miR from the crystal structure 4W5N[15] with a colour gradient from negative (magenta) to positive (blue). The positions in the miR corresponding to the Gs in miR-34a are marked with an arrow in black and red. (b) Variation of amino acid hydrophobicity within 5 Å of the miR from the crystal structure 4W5N[15] is shown with the colour gradient from hydrophilic (purple) to hydrophobic (yellow) based on the Kyte-Doolittle scale[22]. The variation in the electrostatic potential and hydrophobicity along the backbone of miR-34a in the binary complex can potentially explain the increased dispersion in the $^{31}P$ resonance observed in the $^{13}C$-$^{31}P$ TEDOR experiment.
